# Electron transfer proteins in gut bacteria yield metabolites that circulate in the host

**DOI:** 10.1101/2019.12.11.873224

**Authors:** Yuanyuan Liu, Haoqing Chen, William Van Treuren, Bi-Huei Hou, Steven K. Higginbottom, Justin L. Sonnenburg, Dylan Dodd

## Abstract

It has long been known that proteolytic Clostridia obtain their energy by coupling oxidative and reductive pathways for amino acid metabolism – the Stickland reaction^1^. The oxidation of one amino acid is coupled with reduction of another, yielding energy in the former step and re-achieving redox balance with the latter. Here, we find that the gut bacterium, *Clostridium sporogenes* metabolizes amino acids through reductive pathways to produce metabolites that circulate within the host. Measurements *in vitro* indicate that reductive Stickland pathways are coupled to ATP formation, revealing their role in energy capture by gut bacteria. By probing the genetics of *C. sporogenes*, we find that the Rnf complex is involved in reductive amino acid metabolism. Rnf complex mutants are attenuated for growth in the mouse gut, demonstrating the importance of energy capture during reductive metabolism for gut colonization. Our findings reveal that the production of high-abundance molecules by a commensal bacterium within the host gut is linked to an energy yielding redox process.

---

Proteolytic Clostridia are a group of anaerobic bacteria that have the unique ability to grow with amino acids and peptides as their sole energy source. These microbes colonize the gastrointestinal tract of mammals where their metabolic end products accumulate within the gut^2–4^ and are absorbed into circulation mediating important effects on host health^5^. Amino acid metabolites like indolepropionate, 3-(4- hydroxyphenyl)propionate, and indoleacetate are drug-like small molecules that bind host receptors and modulate physiology including intestinal permeability^6, 7^ and immune homeostasis^8, 9^. Amino acid metabolites also mediate syntrophic metabolic interactions among microbes within the gut, with aryl and branched chain fatty acids being required growth factors for several groups of anaerobic gut bacteria^10–12^. Despite their importance to human health and disease, we know very little about the metabolic processes gut bacteria employ to produce such molecules.

## Amino acid metabolism by Clostridium sporogenes

Here we used the gut bacterium, *Clostridium sporogenes* as a model to probe the origin of amino acid metabolites produced within the gut and that circulate in the host. *C. sporogenes* grows rapidly under anaerobic conditions in a defined medium containing a mixture of free amino acids and glucose, achieving a doubling time of ∼37 minutes (**Extended data Fig. 1a**). After 24 h of growth, most amino acids were depleted from the medium (**Extended data Fig. 1b**) and a total of fifteen different metabolites accumulated in culture supernatants (**Extended data Fig. 1c**). Early studies with *C. sporogenes* revealed that it metabolizes amino acids in pairs, coupling the oxidation of one with the reduction of another (the Stickland Reaction)^13^. To assess the degree to which *C. sporogenes* utilizes amino acids as Stickland pairs, we performed a high throughput growth-based assay. Cells were grown in a basal medium containing ten essential amino acids (1 mM each) with higher concentrations (25 mM each) of amino acids or glucose individually or in pairs and growth was monitored spectrophotometrically. The results from these experiments are presented in **Extended data Fig. 2** and summarized in the **Supplementary Text**. Results from these growth-based assays support the following conclusions: 1) Specific pairs of amino acids function synergistically to promote the growth of *C. sporogenes* (Stickland metabolism). 2) Reductive pathway substrates including proline and amino acids that converge on proline (e.g., *trans*-4-hydroxyproline, arginine, citrulline, and ornithine) stimulate growth yields from glucose. 3) *C. sporogenes* utilizes serine in a manner similar to glucose, coupling it to reductive pathway substrates to achieve high growth yields. 4) Proline and *trans*-4-hydroxyproline both stimulated growth yield with branched chain amino acids (valine, isoleucine, and leucine), but this required supplemental acetate presumably for anabolic purposes.

## C. sporogenes produces amino acid metabolites that circulate in the host

To evaluate how *C. sporogenes* influences metabolites that circulate within the host, we colonized germ-free mice with *C. sporogenes* and quantified metabolites in feces, cecal contents, plasma, and urine using stable isotope dilution liquid chromatography – mass spectrometry (LC-MS) (**Supplementary Tables 1-4**). Of the 15 metabolites detected in cultures of *C. sporogenes* (**Extended data Fig. 1c**), 13 were present in the feces of *C. sporogenes* colonized mice one week after colonization (Fig. 1a) demonstrating that *C. sporogenes* produces these metabolites *in vivo*. In plasma, we found four metabolites that were elevated in colonized vs. germ-free mice, which included 5-aminovalerate, phenylpropionate, 3-(4-hydroxyphenyl)propionate, and indolepropionate (Fig. 1a). The remaining nine compounds present in feces, but absent in plasma are likely either excreted in feces or absorbed and metabolized by the host. To confirm that these circulating metabolites arise from bacterial metabolism in the gut, we colonized germ-free mice with *C. sporogenes* and treated separate groups with or without oral antibiotics (Fig. 1b). Following colonization, the four circulating metabolites achieved mean plasma concentrations ranging from ∼3 μM for phenylpropionate to ∼100 μM for indolepropionate (Fig. 1c). After antibiotic treatment, the plasma levels of all four metabolites were reduced to undetectable levels (Fig. 1c), demonstrating their dependence on bacterial metabolism in the gut. In urine, we observed similar trends for 5-aminovalerate, phenylpropionate, and 3-(4-hydroxyphenyl)propionate, with their levels rising after colonization and falling after antibiotic treatment; however, indolepropionate was undetectable in all urine samples (**Supplementary Table 3**). Using a combination of untargeted metabolomics, tandem mass spectrometry (MS/MS), and comparison with authentic standards, we identified high levels of indolepropionylglycine (IPGly) in the urine of *C. sporogenes* colonized mice (**Extended data Fig. 3**) that tracked with plasma levels of indolepropionate (Fig. 1c), suggesting that IPGly is the excreted form of indolepropionate.

**Figure 1.**
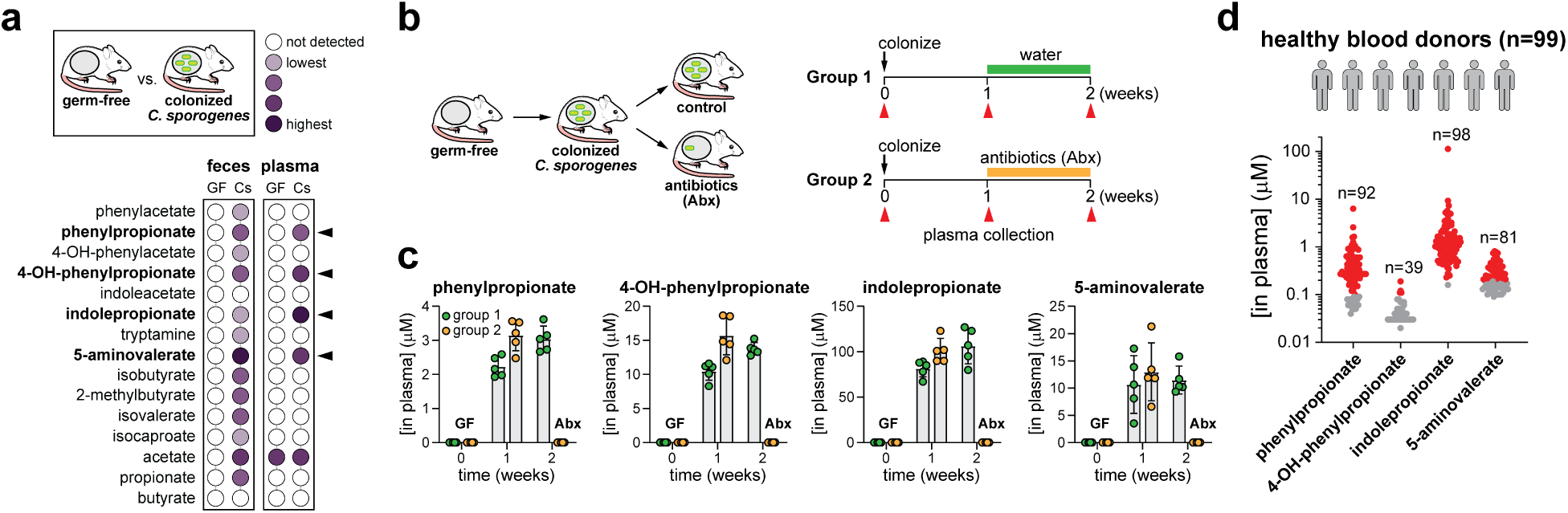
Microbial metabolism within the gut yields metabolites that circulate within the host. A) Fecal and plasma levels of metabolites in germ-free and *C. sporogenes* colonized mice. Data represent means of n = 5 mice. B) Experimental design for detecting metabolites in circulation of gnotobiotic mice before and after antibiotic treatment. C) Quantitation of phenylpropionate, 3-(4- hydroxyphenyl)propionate, indolepropionate, and 5-aminovalerate in plasma of gnotobiotic mice. Abx, antibiotics. D) Quantitation of phenylpropionate, 3-(4- hydroxyphenyl)propionate, indolepropionate, and 5-aminovalerate in plasma of healthy human blood donors (n = 99). Numbers above plots indicate the total number of individuals with detectable metabolite levels. Red dots indicate concentrations within the accurate measurable range of the assay, and gray dots are detected but below the lower limit of quantitation. Concentrations of metabolites detected in mice (A and C) are provided in **Supplementary Tables 1-4** and in humans (D) are provided in **Supplementary Table 5**.

Having identified four *C. sporogenes* dependent circulating metabolites in gnotobiotic mice, we next asked whether these compounds are present in human blood. To test this, we obtained 99 plasma samples from healthy human blood donors and quantified metabolites using LC-MS (**Supplementary Table 5**). Of the four metabolites, indolepropionate was the most prevalent, being detected in the blood of 99% of individuals (Fig. 1d). Consistent with our analysis of gnotobiotic mice (Fig. 1c), indolepropionate concentrations were also the highest, ranging in human plasma from 230 nM to 9.3 μM (Fig. 1d). After repeated analyses, one individual was confirmed to have high plasma indolepropionate levels of 110 μM. Phenylpropionate and 5- aminovalerate were also detected in the majority of healthy blood donors, whereas 3-(4- hydroxyphenyl)propionate levels were lower and present in less than half of individuals (Fig. 1d). Collectively, these findings demonstrate that amino acid metabolism within the gut yields metabolites that circulate within the host. Of these metabolites, indolepropionate is the most concentrated in both gnotobiotic mice and in humans, with rare individuals having high concentrations of circulating indolepropionate.

## Circulating metabolites are products of reductive Stickland metabolism

We next sought to understand the metabolic processes that underlie production of these four circulating metabolites. *C. sporogenes* metabolizes amino acids in pairs, coupling the oxidation of one with the reduction of another via the Stickland Reaction. Oxidative pathways are thought to yield ATP via substrate level phosphorylation, and reductive pathways provide redox balance within the cell (Fig. 2a)^1, 14^. In this context, previous studies have indicated that 5-aminovalerate, phenylpropionate, 3-(4- hydroxyphenyl)propionate, and indolepropionate are end products of reductive Stickland metabolism of Pro, Phe, Tyr, and Trp, respectively^6, 15^.

**Figure 2.**
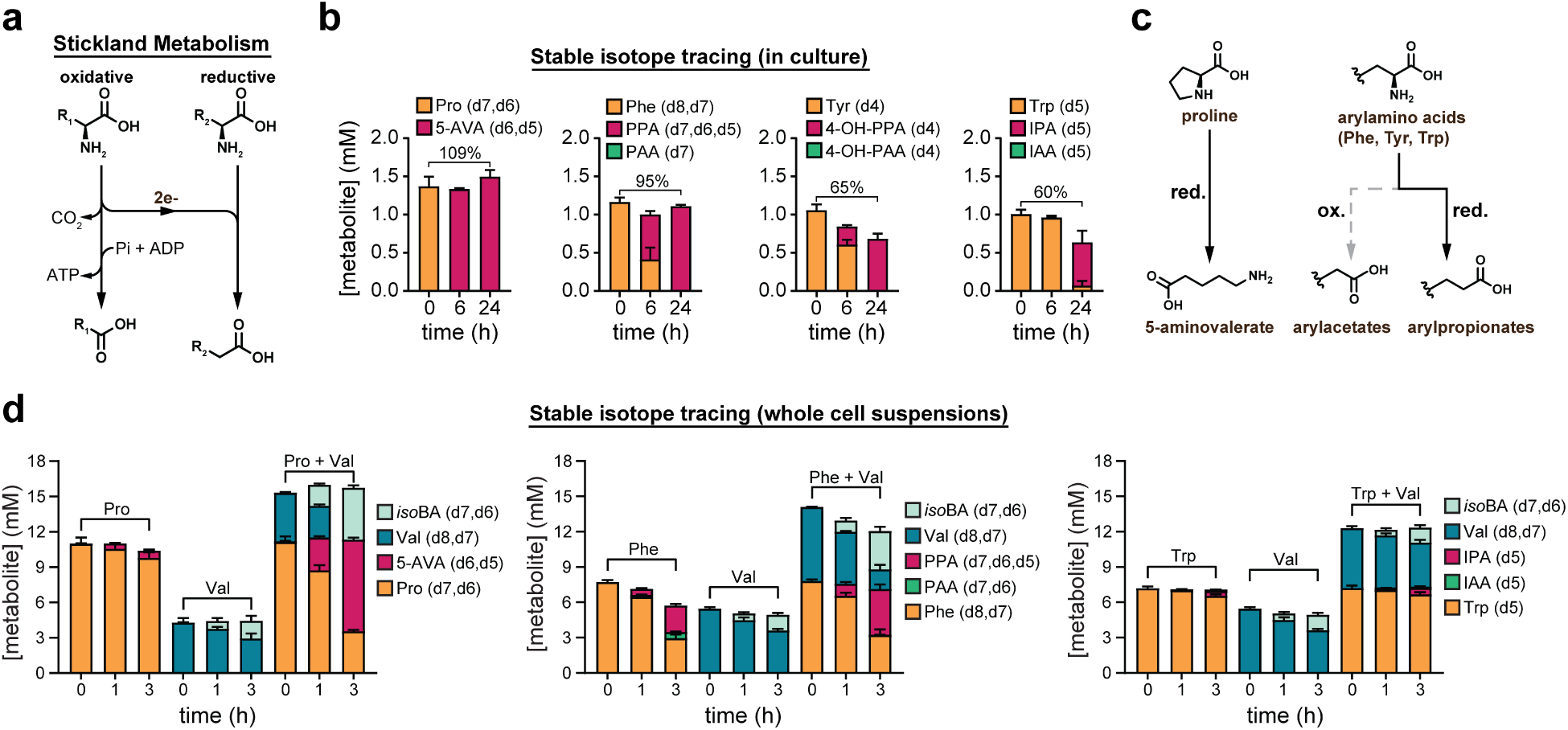
Circulating metabolites are formed from reductive pathways for Stickland metabolism. A) Overview of the Stickland Reaction. Amino acids are metabolized as pairs, with one being oxidized while the other is reduced. The oxidative pathway forms ATP via substrate level phosphorylation and the reductive pathway serves redox balance. B) Stable isotope tracing. *C. sporogenes* was cultured in a synthetic medium containing 20 amino acids where Phe, Tyr, Trp, and Pro were individually substituted by their deuterium isotopologues. Cell-free supernatants (n = 3 per group) were collected at t = 0, 6, 24 h and metabolites were detected by LC-MS. C) End products of Pro, Phe, Tyr, Trp are reductive Stickland metabolites. D) Stable isotope tracing of *C. sporogenes* cell suspensions incubated with oxidative (Val) and reductive (Pro, Phe, Trp) amino acid pairs. 5-AVA, 5-aminovalerate; PPA, phenylpropionate; PAA, phenylacetate; 4-OH-PPA, 3-(4-hydroxyphenyl)propionate; 4- OH-PAA, 4-hydroxyphenylacetate; IPA, indolepropionate; IAA, indoleacetate; *iso*BA, isobutyrate.

To test whether these four circulating metabolites arise from metabolism of their cognate amino acids, we performed stable isotope tracing experiments. *C. sporogenes* was cultured in a defined medium containing all 20 proteinogenic amino acids (1 mM each) and glucose (10 mM) where Pro, Phe, Tyr, or Trp were individually substituted by their deuterium isotopologues (Pro-d7, Phe-d8, Tyr-d4, or Trp-d5) and metabolites were measured by LC-MS during growth. Under these conditions, cells reached maximal optical densities of ∼0.5 by 16 h. Labeled Pro, Phe, and Tyr were completely consumed by 24 h, being converted to labeled derivatives (5-aminovalerate-d6,d5; phenylpropionate-d7,d6,d5; and 3-(4-hydroxyphenyl)propionate-d4) with yields ranging from 65%-109% (Fig. 2b). Trp metabolism was comparably slower, but ∼93% of Trp was consumed by 24 h, being converted to indolepropionate-d5 with a yield of 60% (Fig. 2b). Despite pathways existing in *C. sporogenes* for the oxidative metabolism of Phe, Tyr, and Trp^6, 16^, under these experimental conditions only reductive products of these amino acids were detected (Fig. 2c). These findings indicate that *C. sporogenes* cells convert the majority of these amino acids to reduced products, rather than using them for biosynthetic purposes such as protein synthesis. In cultures where labeled Phe, Tyr, and Trp were provided, no unlabeled phenylpropionate, 3-(4- hydroxyphenyl)propionate, or indolepropionate was detected (**Extended data Fig. 4a-c**) suggesting that these metabolites arise from catabolism of their cognate amino acids, not through *de novo* biosynthetic pathways from other precursors. 5-aminovalerate was an exception, however, with its unlabeled product accumulating in cultures where labeled Pro was added (**Extended data Fig. 4d**), which we found reflected the capacity of cells to convert Arg in the medium to proline and 5-aminovalerate (**Extended data Fig. 4e**)^17^.

During Stickland metabolism, the oxidation of one amino acid is coupled to the reduction of another, resulting in stimulatory effects of amino acid pairs^1^. Next, we sought to test whether production of the reductive metabolites, 5-aminovalerate, phenylpropionate, and indolepropionate is stimulated by provision of an oxidative amino acid partner. Cell suspensions of *C. sporogenes* were incubated with isotopically labeled amino acids (Pro, Phe, Trp) in the presence or absence of an oxidative Stickland amino acid (Val) and metabolism was monitored by LC-MS. Whereas cells incubated with Pro or Val alone showed limited substrate consumption and product accumulation (Fig. 2d), cells incubated with both Pro and Val consumed most of the Pro and all the Val by 3h, converting Pro to 5-aminovalerate and Val to isobutyrate (Fig. 2d). Similar results were seen with the Phe/Val combination (Fig. 2d), however the Trp/Val combination yielded a more subtle pattern, where indolepropionate production was low (Fig. 2d), but significantly higher when Val was added (**Extended data Fig. 5**). Collectively, these findings demonstrate that 5-aminovalerate, phenylpropionate, 3-(4- hydroxyphenyl)propionate, and indolepropionate are reductive pathway products of coupled Stickland reactions by gut bacteria.

## Reductive Stickland metabolism is coupled to ATP formation

During Stickland metabolism of amino acids, oxidative pathways are thought to yield ATP via substrate level phosphorylation, and reductive pathways provide redox balance within the cell^1, 14^. However, anaerobic bacteria can also use reductive metabolism to fuel anaerobic respiration, providing an additional mechanism to obtain energy. Examples include fumarate reduction by *Bacteroides fragilis* (involved in carbohydrate metabolism)^18^, caffeate respiration by *Acetobacterium woodii*^19^ and cinnamate reduction by *C. sporogenes*^20^ (involved in metabolism of plant secondary metabolites), and reductive acetogenesis by *C. ljungahlii*^21^. We reasoned that reductive pathways for amino acid metabolism via the Stickland reaction might also serve a role in energy capture for bacteria within the gut. A previous study demonstrated that Pro reduction in *C. sporogenes* is linked to vectorial transmembrane proton translocation^22^, however whether this proton gradient is coupled to ATP synthesis is unknown. Furthermore, it is not known whether additional pathways for reductive amino acid metabolism are involved in energy capture. Therefore, we analyzed the metabolic gene clusters encoding reductive amino acid pathways with an eye towards identifying potential mechanisms for energy capture.

Inspection of the gene clusters encoding reductive amino acid pathways revealed that each pathway converges on a step involving a multi-subunit electron transfer protein (Fig. 3a). The aryl amino acid reductive gene cluster encodes an acyl-CoA dehydrogenase with two electron transfer factors (EtfAB) sharing 40-60% amino acid identity with the butyryl-CoA dehydrogenase-EtfAB (Bcd) complex from *Clostridium kluyverii* involved in ethanol fermentation^23^ (Fig. 3b). Bcd catalyzes flavin based electron bifurcation where the energetically downhill reduction of crotonyl-CoA by NADH drives the energetically uphill reduction of ferredoxin by NADH^24, 25^. In so doing, the cell has low-potential reduced ferredoxin at its disposal to drive ion translocation through the Rnf complex with NAD^+^ as an electron acceptor^26^. The gene cluster for reductive leucine metabolism, present in *C. sporogenes*^27^, also encodes a homolog of the Bcd complex (Fig. 3b). The proline reductive gene cluster encodes a different electron transfer protein involving the enzyme proline reductase (Fig. 3c). However, a link to energy conservation for proline reductase has not been established. Thus, these metabolic gene clusters suggest that reductive Stickland pathways may be linked to ATP synthesis in the cell.

**Figure 3.**
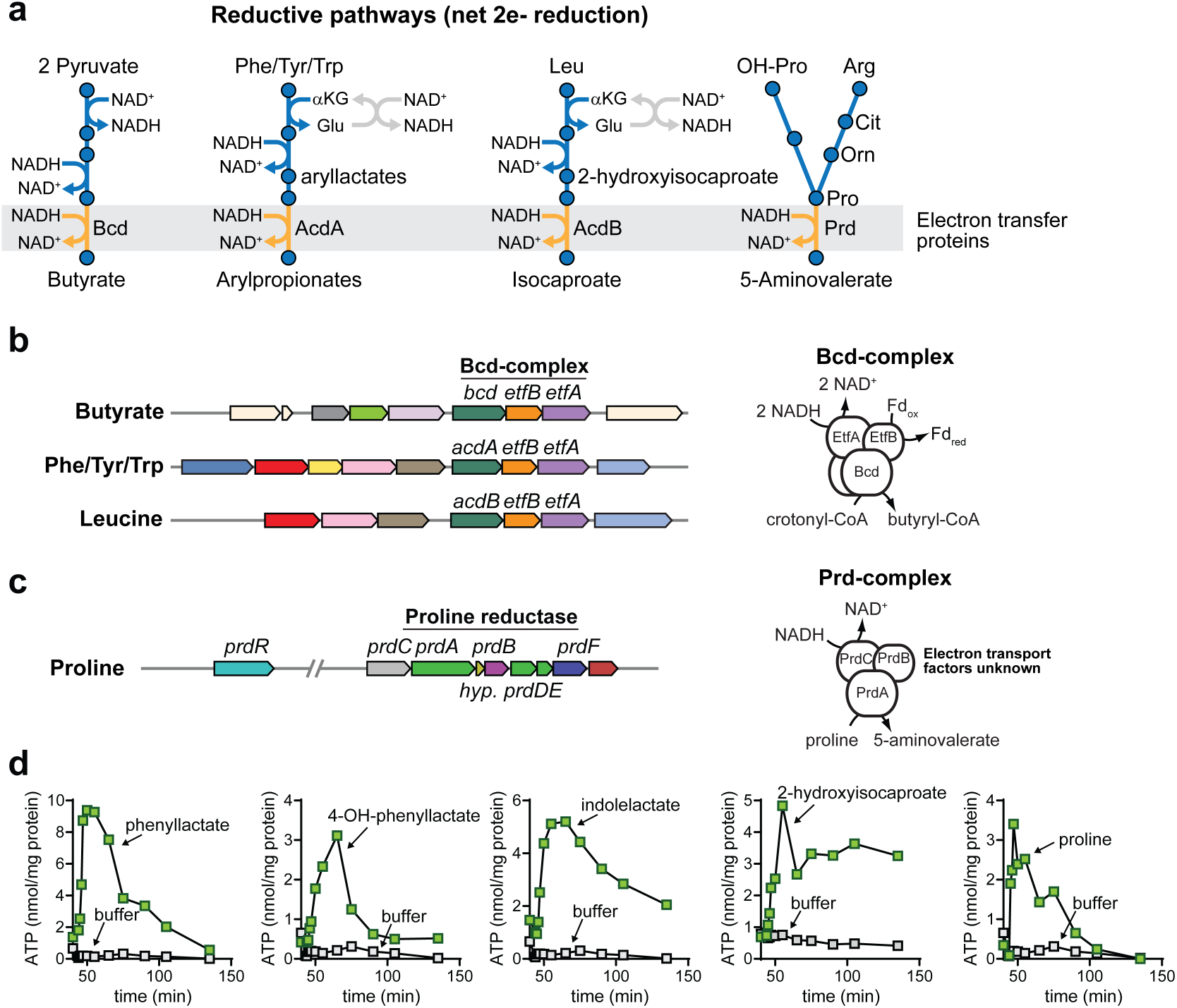
Reductive metabolism is coupled to ATP formation. A) Pathways for reductive Stickland metabolism catalyze net 2 electron reductions and each converges on an electron transfer protein. B) Gene clusters for reductive metabolism of Phe, Tyr, Trp, and Leu share homologs of the electron bifurcating butyryl-CoA dehydrogenase complex (Bcd) which produces reduced ferredoxin. C) The proline metabolism gene cluster encodes components of the proline reductase enzymes which is implicated in proline-dependent extracellular proton transport. D) Reductive metabolism of Phe, Tyr, Trp, Leu, and Pro are coupled to ATP formation in *C. sporogenes* resting cell suspensions. For Phe, Tyr, and Trp, intermediates in the reductive pathways (e.g., phenyllactate, 3-(4-hydroxyphenyl)lactate, and indolelactate) were used. For D, experiments were repeated independently three times and representative data are shown.

To test this experimentally, we established a whole cell-based ATP assay using resting late log-phase cell cultures of *C. sporogenes*. Following incubation with substrate, cells were quenched and permeabilized with DMSO, then ATP levels were detected by luminescence and normalized to total cellular protein levels. We validated our assay by analyzing ATP formation from *trans*-cinnamate, a plant derived substrate previously thought to be involved in reductive phenylalanine metabolism (**Extended data Fig. 6, Supplementary Text**). Because Phe, Tyr, Trp, and Leu may also be metabolized down oxidative pathways, we supplied cells with pathway intermediates (e.g., phenyllactate, 3-(4-hydroxyphenyl)lactate, indolelactate, 2-hydroxyisocaproate) specific to reductive pathways. When Pro or reductive pathway intermediates for Phe, Tyr, Trp, or Leu were added to cells, ATP levels rose rapidly, plateaued, and then returned to resting levels (Fig. 3e). Because reductive arylamino acid metabolism (Phe, Tyr, Trp) operates through a shared pathway, we chose to further explore ATP formation using Phe metabolism as a representative for this pathway. These analyses revealed that D-phenyllactate (and not its L-stereoisomer) supported ATP formation (**Extended data Fig. 7a**), being converted almost quantitatively to the reductive pathway product, phenylpropionate (**Extended data Fig. 7b**). Metabolic products (**Extended data Fig. 7c**) of reductive or oxidative pathways failed to elicit ATP formation (**Extended data Fig. 7d-e**). Genetic disruption of acyl-CoA dehydrogenase (AcdA) abolished ATP formation from DL-phenyllactate, indicating that this enzyme is critical for ATP formation during reductive Phe metabolism (**Extended data Fig. 7f**). Similar results were seen for metabolism of the Tyr and Trp metabolic intermediates (DL-4-hydroxyphenyllactate and DL-indolelactate), with the *acdA* mutant being deficient in ATP formation from these substrates (**Extended data Fig. 7g-h**). Pre-treatment of cells with the protonophore, 3,3′,4′,5-tetrachlorosalicylanilide (TCS) uncoupled DL- phenyllactate and proline metabolism from ATP formation (**Extended data Fig. 8**), indicating that a proton gradient is involved in ATP formation. Taken together, these experiments reveal that pathways for reductive Stickland metabolism are linked to ATP formation and involve the generation of a proton motive force.

## The Rnf complex is important for reductive metabolism and competition within the gut

Having demonstrated that reductive metabolism is coupled to ATP formation, we next asked whether the Rnf complex might be an important coupling site. We identified a gene cluster which encodes the Rnf complex in *C. sporogenes* (Fig. 4a) and targeted genes encoding two separate subunits (*rnfB* and *rnfE*) for disruption (Fig. 4b). These mutants suffered a growth defect when grown in minimal medium containing 10 amino acids (Fig. 4c), yet this phenotype could be suppressed by the addition of glucose to the culture medium (Fig. 4c). These data suggest that the growth phenotype in *rnfB* and *rnfE* mutants is characterized by defective amino acid metabolism. Metabolic profiling of culture supernatants revealed that the *rnfB* mutant was defective in production of the reductive pathway metabolites 5-aminovalerate, phenylpropionate, 3-(4- hydroxyphenyl)propionate, and indolepropionate, suggesting an impairment in reductive Stickland metabolism (Fig. 4d**, Extended data Fig. 9**). Thus while the Rnf complex clearly plays a role in amino acid metabolism, it is unclear whether the mutant phenotype is due solely to diminished ATP formation during reductive metabolism, or a combined loss of regenerative functions that drive forward oxidative pathways.

**Figure 4.**
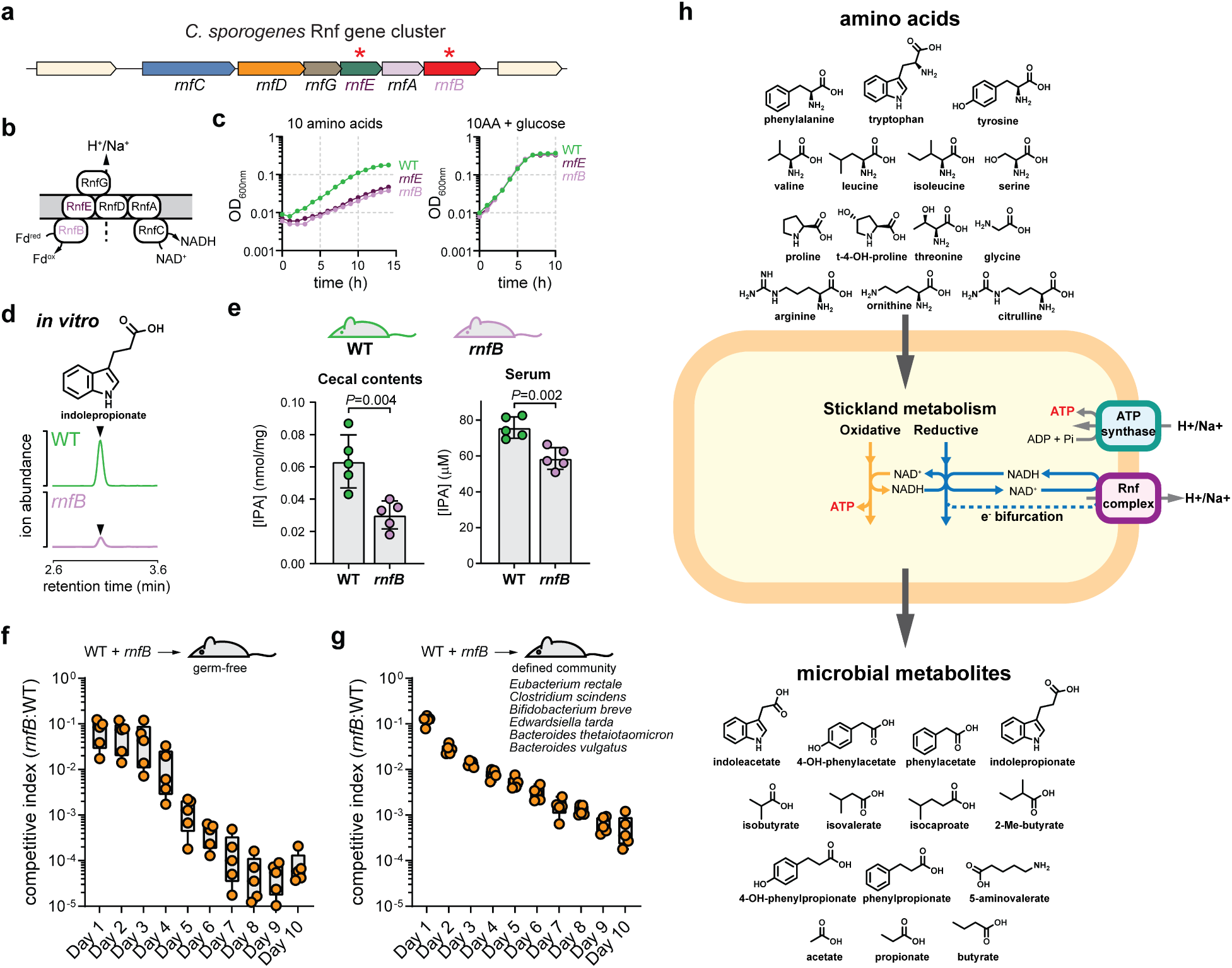
The Rnf complex is important for amino acid metabolism and is a key fitness determinant in vivo. A) The *C. sporogenes* Rnf complex gene cluster. B) Model for the membrane bound Rnf complex which generates a proton or sodium motive force by shuttling electrons from reduced ferredoxin to NAD^+^. C) The *rnfB* and *rnfE* mutants suffer a growth defect in amino acid medium that can be restored by addition of glucose. D) The *rnfB* mutant produces less indolepropionate during growth in defined medium with 10 amino acids. E) Mice mono-colonized with the *rnfB* mutant accumulate less indolepropionate in feces and in plasma. F) The *rnfB* mutant exhibits an *in vivo* competition defect in gnotobiotic mice in the context of mono-colonization and G) a more complex defined microbial community. H) Revised model for Stickland metabolism and its contribution to gut bacterial derived metabolites. For C-D, experiments were repeated independently three times and representative data are shown. For E-G, Box and whisker plots show median values, 25th–75th percentiles and range for *n* = 5 biological replicates. IPA, indolepropionate.

Our findings suggest that disruption of the Rnf complex leads to altered amino acid metabolism *in vitro*. Next, we asked whether energy capture through the Rnf complex is important for production of reduced metabolites that circulate within the host. To test this, we colonized gnotobiotic mice with either WT *C. sporogenes* or its *rnfB* mutant and detected metabolites in cecal contents and plasma by LC-MS. By comparison to WT-colonized mice, *rnfB*-colonized mice had significantly lower levels of indolepropionate both in the cecal contents and the plasma (Fig. 4e), demonstrating a role for the Rnf complex in production of microbiota-derived metabolites that circulate within the host. To test whether energy capture involving the Rnf complex is important for competition within the mammalian gut, germ-free mice were colonized with a 1:1 mixture of WT and *rnfB* mutant cells and the relative proportions of the two strains in feces were determined by Q-PCR over time. The *rnfB* mutant was rapidly outcompeted by the WT strain, becoming nearly undetectable in feces by 7 days post colonization (Fig. 4f). This finding was recapitulated in the presence of a more complex stably colonized microbial community revealing the Rnf complex as a key fitness determinant for *C. Sporogenes* during colonization and persistence in the gastrointestinal tract (Fig. 4g**, Extended data Fig. 10**). While our results suggest the Rnf complex is important for ATP formation, the loss of a proton gradient that Rnf presumably creates, could influence additional physiologic processes such as membrane transport and chemotaxis.

Intriguingly, the Rnf genes were previously identified as *in vivo* fitness determinants for *Bacteroides* colonization in the mouse gut^28, 29^ yet were dispensable for growth in defined medium^28^. This suggests that the Rnf complex, which is encoded by phylogenetically diverse human gut bacteria (**Extended data Fig. 11**), may serve distinct physiological roles across taxa.

## Conclusion

Our results show that *C. sporogenes* ferments amino acids through coupled oxidative and reductive pathways (the Stickland reaction), to produce metabolites that accumulate in the intestinal lumen and that circulate within the host (Fig. 4h). In addition to producing ATP via substrate level phosphorylation, reductive Stickland pathways provide additional energy likely via electron transport phosphorylation. Acyl-CoA dehydrogenases (Bcd/AcdA/AcdB) capture this energy via flavin-based electron bifurcation, where potential energy is carried by reduced ferredoxin which then drives extracellular proton (or sodium ion) transport via the Rnf complex. Homologs of the electron transfer proteins described in this study are present in bacteria from 7 unique Phyla and 38 different families (**Supplementary Table 6**). They are also prevalent within human fecal metagenomic datasets (**Extended data Fig. 12**), suggesting these mechanisms for energy capture are widespread within the gut. By reevaluating ATP economy in the cell (considerations in **Supplementary Table 7**), reductive pathways can account for ∼40% of ATP from Stickland metabolism. This revises our understanding of how amino acid degrading bacteria capture their energy and provides fundamental insights into the origin of metabolites that circulate within the human body.

## Methods

### Bacterial strains and culture conditions

*Clostridium sporogenes* ATCC 15579 was obtained from the American Type Culture Collection (ATCC). It was routinely cultured in reinforced clostridial medium (RCM) at 37 °C in a Coy type B anaerobic chamber under an atmosphere of 5% hydrogen, 10% carbon dioxide, and 85% nitrogen. All media and plasticware were pre-reduced in the anaerobic chamber for at least 24 hours before use. For growth measurements in defined medium, we used a previously described defined minimal medium referred to as standard amino acid complete medium (SACC)^15^. The *Escherichia coli* CA434 (HB101/pRK24) conjugation host was routinely cultured at 30 °C in LB broth supplemented with tetracycline (12 μg mL^-1^) to ensure maintenance of the pRK24 plasmid. *E. coli* TG1 was used for routine cloning. Chloramphenicol (25 μg mL^-1^) was used for selection of pMTL007C-E2 plasmids in *E. coli*. Organisms used in the defined community colonization included: *Clostridium sporogenes* ATCC 15579, *Clostridium scindens* ATCC 35704, *Eubacterium rectale* ATCC 33656, *Bifidobacterium breve* UCC2003, *Edwardsiella tarda* ATCC 23685, *Bacteroides thetaiotaomicron* VPI-5482, and *Bacteroides vulgatus* ATCC 8482.

### Plasmid construction and cloning

For specific gene disruptions, we used the Intron targeting and design tool on the ClosTron website (http://www.clostron.com/clostron2.php) using the Perutka algorithm. The intron contained within the pMTL007C-E2 plasmid was retargeted to the specific sites listed in **Supplementary Table 8** and were synthesized by DNA 2.0 (Menlo Park, CA). Sequencing primers were synthesized by IDT DNA Technologies and are listed in Supplementary Table 8.

### Mutant generation using ClosTron

Intron-retargeted ClosTron plasmid DNA was introduced into *C. sporogenes* by conjugation as described previously^6^. Transconjugants were selected on RCM agar supplemented with D-cycloserine (250 μg mL^−1^) and thiamphenicol (15 μg mL^−1^), re-streaked for purity, and individual well-isolated colonies were inoculated into RCM broth (without agar) containing the same antibiotics. After overnight culture, the cells were diluted 10-fold, then 100 μL was spread on an RCM agar plate containing erythromycin (5 μg mL^−1^). These colonies were picked, re-streaked onto TYG agar plates supplemented with erythromycin and well-isolated colonies were inoculated into TYG broth supplemented with erythromycin. Genomic DNA was isolated from candidate clones using the DNeasy Blood and Tissue Kit from Qiagen with a lysozyme pre-treatment step, and this DNA was used as a template for PCR using gene-specific primers. Primer sets were designed to produce a ∼600-bp product for the wild-type and ∼2,800-bp product for the mutant containing the intron. When verified mutants were streaked onto TYG plates containing thiamphenicol, no growth was observed, indicating that the pMTL007C-E2 plasmid had been lost.

### Monitoring bacteria growth in batch culture and analysis of metabolites in culture supernatants by GC-MS and LC-MS

To characterize the growth of *C. sporogenes* with amino acids (**Extended data Fig. 1a**), cells were cultured in SACC medium supplemented with 23 amino acids including the 20 proteinogenic amino acids and L-citrulline, L-ornithine, and *trans*-4-hydroxyproline (1 mM each) and glucose (1 mM). Wild-type *C. sporogenes* was streaked from an anaerobic glycerol stock onto RCM agar plates and incubated for ∼24 h at 37 °C. Individual well-isolated colonies were then inoculated into 5 mL of SACC (23 AA + glucose) medium and cultured for 24 h at 37 °C. Optical densities at 600 nm (OD600 nm) were recorded for cultures, then cells were subcultured in 5 mL of SACC (23 AA + glucose) at a starting OD of 0.01 in butyl rubber stoppered anaerobic Balch-type tubes (ChemGlass). Cultures were incubated outside of the anaerobic chamber in a water bath set to 37 °C, and optical density measurements were made hourly using a GENESYS 30 spectrophotometer from Thermo Scientific. Samples (1 mL) were collected at 0 and 24 h, centrifuged at 5,000*g* for 10 min at 4 °C, then supernatants (500 μL) were collected and stored at −20 °C until analysis by GC-MS or LC-MS.

To characterize the growth phenotype of *rnfB* and *rnfE* mutants (Fig. 4c), cells were cultured as above with the following minor modifications. The medium used was SACC medium with 10 essential amino acids (1 mM each), with or without glucose (10 mM). Mutant cells were first streaked from anaerobic glycerol stocks onto RCM agar plates with erythromycin (5 μg mL^−1^), to confirm they carried the Clostron insertion, and all subsequent growth was performed without supplemental antibiotics.

#### Amino acid analysis in culture supernatants (corresponding to Extended data Fig. 1b)

Culture supernatant was diluted 80-fold with LC-MS grade water, then 50 μL of samples or appropriately diluted standards (Sigma AAS18 supplemented with Trp, Asn, Gln, ornithine, *trans*-4-hydroxyproline, and 5-aminovalerate) were combined with 10 μL internal standards (L-Arginine:HCl (Guanido-^15^N_2_), DL-Valine (D8), L-Methionine (Methyl-D3), L-Phenylalanine (D8), L-Leucine (D10), and L-Tyrosine (D7), 10 μM each) in 96-well V-bottom plates from USA scientific. Then 25 μL of 0.1 M sodium tetraborate and 50 μL of dansyl chloride (25 mM in 100% acetonitrile) were added. The solution was mixed five times by pipetting, then incubated at room temperature in the dark for 15 min, mixed by pipetting five times, then incubated for an additional 15 min. The reaction was quenched by the addition of 50 μL 0.5% formic acid, and the plate was centrifuged at 5,000*g* for 15 min at room temperature. One hundred and twenty-five microliters were then transferred to a Multiscreen Solvinert Filter Plate 0.45 μM (Millipore) placed on top of a 96-well 0.5 mL polypropylene microplate (Agilent) which was then centrifuged at 5,000*g* for 5 min at room temperature. A cap mat was placed on the plate, and samples were analyzed by LC-MS using an Agilent 1290 Infinity II UPLC equipped with a Waters BEH C18 1.7-μm particle size C18 column (2.1 × 100 mm) and detected using an Agilent 6545XT Q-TOF equipped with a dual jet stream electrospray ionization source operating under extended dynamic range (EDR 1700 m/z) in positive ionization mode. Eluent A consisted of 0.1% formic acid (v/v) in water and eluent B consisted of 0.1% formic acid (v/v) in 100% acetonitrile. Five microliters of samples were injected via refrigerated autosampler into mobile phase and chromatographic separation at 40 °C was achieved at a flow rate of 0.5 mL min^−1^ using a linear gradient of 5–60% eluent B over 10.25 min. Eluent B was brought to 95% over 2.25 min, held at 95% for 2.5 min, brought to 5% over 0.1 min, and the column was then re-equilibrated at 5% eluent B for 2.9 min. Electrospray ionization parameters included a gas temperature of 325 °C, drying gas flow of 10 L min^-1^, nebulizer pressure of 20 psi, sheath gas temperature of 400 °C and flow of 12 L min^-1^, and capillary voltage of 4000 V. MS1 spectra were collected in profile mode, and peak assignments in samples were made based on comparisons of retention times and accurate masses from authentic standards. Amino acids were quantified by isotope-dilution mass spectrometry using L-Phenylalanine (D8) as a suitable internal standard for most analytes.

#### Aromatic fatty acids and amines by LC-MS (corresponding to Extended data Fig. 1c, Extended data Fig. 6, Fig. 4d)

Culture supernatants or appropriately diluted standards (phenylacetate, phenylpropionate, 4-hydroxyphenylacetate, 3-(4- hydroxyphenyl)propionate, indoleacetate, indolepropionate, tryptamine, 5- aminovalerate) (100 μL) were combined with 450 μL of methanol followed by incubation at room temperature for 5 min and centrifugation at 13,000*g* for 5 min at room temperature. Subsequently, 200 μL of supernatant was transferred to wells of a 96-well deep block plate and evaporated to dryness using a Biotage Turbovap. Then 200 μL of resuspension solvent (0.1% formic acid (v/v) in water) containing internal standards (100 μM *trans*-cinnamic acid (D7) and 50 μM indoleacetic acid (D5)) was added, compounds were dissolved by vortexing for ∼30 s and samples were clarified by filtration through a Multiscreen Solvinert Filter Plate 0.45 μM (Millipore) placed on top of a 96-well 0.5 mL polypropylene microplate (Agilent) by centrifugation at 5,000*g* for 5 min at room temperature. A cap mat was placed on the plate, and samples were analyzed by LC-MS using an Agilent 1290 Infinity II UPLC equipped with a Waters BEH C18 1.7- μm particle size C18 column (2.1 × 100 mm) and detected using an Agilent 6545XT Q- TOF equipped with a dual jet stream electrospray ionization source operating under extended dynamic range (EDR 1700 m/z) in negative ionization mode. Eluent A consisted of 6.5 mM ammonium bicarbonate in water and eluent B consisted of 6.5 mM ammonium bicarbonate in 100% methanol. Five microliters of samples were injected via refrigerated autosampler into mobile phase and chromatographic separation was achieved at a flow rate of 0.35 mL min^−1^ using a linear gradient of 0.5–70% eluent B over 4 min. Eluent B was brought to 98% over 0.5 min, held at 98% for 0.9 min, then brought to 0.5% over 0.2 min and the column was re-equilibrated at 0.5% eluent B for 3.4 min. Peak assignments in samples were made based on comparisons of retention times and accurate mass-to-charge ratios from authentic standards analyzed under identical conditions. Electrospray ionization parameters included a gas temperature of 250 °C, drying gas flow of 6 L min^-1^, nebulizer pressure of 30 psi, sheath gas temperature of 200 °C and flow of 11 L min^-1^, capillary voltage of 4000 V, and nozzle voltage of 1400 V. MS1 spectra were collected in centroid mode, and peak assignments in samples were made based on comparisons of retention times and accurate masses from authentic standards. Aromatic fatty acids were quantified by isotope-dilution mass spectrometry using *trans*-cinnamic acid (D7) or indoleacetic acid (D5) as internal standards.

#### Short chain fatty acids by GC-MS (corresponding to Extended data Fig. 1c)

Culture supernatants (50 μL) were acidified with 6 M HCl (50 μL), then diluted in LC-MS grade water (150 μL). Organic acids were then extracted with one volume of diethyl ether (250 μL). An aliquot (95 μL) of the organic layer was transferred to a new sealed vial, combined with *N-tert*-butyldimethylsilyl-*N*-methyltrifluoroacetamide (MTBSTFA, 5 μl) and derivatized at room temperature for 48 h. One microliter was then injected onto an Agilent 7820A gas chromatograph equipped with an Agilent VF-5HT column (30 m x 0.25 mm x 0.1 μm) coupled to an Agilent 5977B mass spectrometer. The inlet was set to 280 °C, and injection occurred in split mode with a ratio of 10:1. Oven conditions were as follows: 50 °C for 2 min, ramped to 70 °C at 10 °C min^-1^, then ramped to 85 °C at 3 °C min^-1^, then ramped to 290 °C at 30 °C min^-1^. The mass spectrometer transfer line was set to 250 °C, and the quadrupole was set at 150 °C. MS data was collected under scan mode from 50 – 300 m/z with a 3 min solvent delay.

### Wild-type C. sporogenes mono-colonization in gnotobiotic mice and analysis of metabolites in body fluids (corresponding to Fig. 1a,c, Supplementary Tables 1-4).

Mouse experiments were performed on gnotobiotic Swiss Webster germ-free mice originally obtained from Taconic Biosciences maintained in aseptic isolators. Animal experiments were performed following a protocol approved by the Stanford University Administrative Panel on Laboratory Animal Care. Mice (female, 9 weeks of age, n = 5 per group, two groups) were colonized with wild-type *C. sporogenes* by oral gavage (300 μL, ∼1 × 10^8^ CFU) and were maintained on standard chow (LabDiet 5k67). After one week, group 1 mice were maintained on normal drinking water, while group 2 mice were switched to water containing antibiotics (0.5 mg/mL Vancomycin and 1 mg/mL Metronidazole) for another week before euthanization. Blood, urine and feces were collected at 3 time points: before gavage with bacteria, 1 week and 2 weeks after colonization. Cecal contents were harvested after euthanization. Blood collected from the facial vein was mixed with EDTA (∼15 mM EDTA, final) and centrifuged at 1,500*g* for 15 min at 10 °C. Plasma was transferred to a new tube and stored at −80 °C until LC-MS was performed. Fresh urine, fecal pellets, and cecal contents were collected and stored at −80 °C until LC-MS was performed.

#### Sample preparation for metabolite analysis in blood, urine, feces, and cecal contents

Mice urine samples (2.5 μL) were diluted with 7.5 μL LC-MS water. Blood (10 μL) and diluted urine (10 μL) samples were added to a 96-well V-bottom plate containing internal standard mixture (tryptamine (D4), indoleacetic acid (D5), phenylacetic acid (D5), phenylpropionic acid (D9), phenylalanine (D5), hippuric acid (D2), 0.2 mM each, creatinine (D3), acetic acid (^13^C_2_), isovaleric acid (D2), 1 mM each, 10 μL) and 180 μL methanol. The solution was mixed five times by pipetting, then the plate was centrifuged at 5,000*g* for 15 min at 4 °C. Supernatant was collected for the following sample preparations, either derivatization or direct dilution, before subjecting to LC-MS analysis.

Mice cecal contents and fecal samples (40(±4) mg) were weighed in microcentrifuge tubes (Fisherbrand cat. # 02-682-558) containing glass beads (Sigma cat. # G1145, 150-212 μm, 40(±4) mg). LC-MS water (40 μL), internal standard mixture (tryptamine (D4), indoleacetic acid (D5), phenylacetic acid (D5), phenylpropionic acid (D9), phenylalanine (D5), 0.2 mM, acetic acid (^13^C_2_) and isovaleric acid (D2), 1 mM each, 80 μL), and methanol (1440 μL) were added to the tubes. The samples were homogenized with a mixer mill (RETSCH MM400) at 4 °C, 25/s, for 30 min, then centrifuged at 13,000*g* for 5 min at 4 °C. Supernatants were collected for the following sample preparations, either derivatization or direct dilution, before subjecting to LC-MS analysis.

#### Derivatization method for short chain fatty acids

Thirty microliters of supernatant was transferred to a 96-well V-bottom plates and mixed with 15 μL 3- nitrophenylhydrazine (200 mM in 50% acetonitrile) and 15 μL *N*-(3- dimethylaminopropyl)-*N’*-ethylcarbodiimide (120 mM in 6% pyridine). The plates were sealed with a plastic sealing mat (Thermo Fisher Scientific cat. # AB-0566) and incubated at 40 °C, 900 rpm in a thermomixer for 30 min to derivatize the short chain fatty acids. Eight microliters of the reaction mixture were quenched with 192 μL 0.02% formic acid in 10% acetonitrile/water.

#### Direct dilution method for other metabolites

Thirty microliters of supernatant were directly transferred to a 96-well V-bottom plates and diluted with LC-MS grade water (90 μL) for LC-MS analysis.

#### LC-MS conditions for detection and quantitation of metabolites

The samples were analyzed using the same instrument and column with the following methods. (i) Derivatized acetate, propionate, butyrate, isobutyrate, 2-methylbutyrate, isovalerate, isocaproate, 5-aminovaleric acid, indoleacetate, indolepropionate, phenylacetate, phenylpropionate, 3-(4-hydroxyphenyl)acetate, 3-(4-hydroxyphenyl)propionate, hippurate, phenylacetylglycine, phenylpropionylglycine, 4-hydroxyhippurate, (3-(4- hydroxyphenyl)propanoyl)glycine, and 4-hydroxyphenylacetylglycine were analyzed under negative ionization mode. Eluent A consisted of 0.1% formic acid (v/v) in water and eluent B consisted of 0.1% formic acid (v/v) in 100% methanol. Samples (10 μL) were injected via refrigerated autosampler into mobile phase and separation was achieved with Waters BEH C18 1.7-μm particle size C18 columns (2.1 × 100 mm) at 50 °C using dual binary pumps with alternating column regeneration. The analytical pump applied the following gradient at a flow rate of 0.3 mL min^−1^: 0−11 min, 15% to 65% B, 11−11.1 min, 65% to 15% B, 11.1 min−12 min, 15% B. The regeneration pump applied the following gradient at a flow rate of 0.2 mL min^−1^: 0−0.1 min, 15% to 99.9% B, 0.1−6 min, 99.9% B, 6−9 min, 99.9% to 15% B, 9−12 min, 15% B. Electrospray ionization parameters included a gas temperature of 300 °C, drying gas flow of 6 L min^-1^, nebulizer pressure of 30 psi, sheath gas temperature of 300 °C and flow of 11 L min^-1^, capillary voltage of 4000 V, and fragmentor voltage of 140 V. (ii) Underivatized creatinine and tryptamine were analyzed under positive ionization mode. Eluent A consisted of 0.1% formic acid (v/v) in water and eluent B consisted of 0.1% formic acid (v/v) in 100% methanol. Samples (0.5 μL) were injected via refrigerated autosampler into mobile phase and separation was achieved with Waters BEH C18 1.7-μm particle size C18 columns (2.1 × 100 mm) at 25 °C using dual binary pumps with alternating column regeneration. The analytical pump applied the following gradient at a flow rate of 0.3 mL min^−1^: 0−4 min, 1% to 70% B, 4−4.01 min, 70% to 1% B, 4.01−5 min, 1% B. The regeneration pump applied the following gradient at a flow rate of 0.3 mL min^−1^: 0−0.1 min, 1% to 98% B, 0.1−3 min, 98% B, 3−3.1 min, 98% to 1% B, 3.1−5 min, 1% B. Electrospray ionization parameters included a gas temperature of 250 °C, drying gas flow of 6 L min^-1^, nebulizer pressure of 30 psi, sheath gas temperature of 250 °C and flow of 11 L min^-1^, capillary voltage of 4000 V, and fragmentor voltage of 70 V.

Selection of internal standards (ISTD) for LC-MS quantitation were as follows: isovaleric acid (D2) as ISTD for butyrate, isobutyrate, 2-methylbutyrate, isovalerate, and isocaproate. Indoleacetic acid (D5) as ISTD for indoleacetate, indolepropionate, 4- hydroxyhippurate, (3-(4-hydroxyphenyl)propanoyl)glycine, and 4- hydroxyphenylacetylglycine. Phenylacetic acid (D5) as ISTD for phenylacetate. Phenylpropionic acid (D9) as ISTD for phenylpropionate, 4-OH-phenylpropionate, 4-OH- phenylacetate, and phenylacetylglycine. Acetic acid (^13^C_2_) as ISTD for acetate, propionate, and 5-aminovaleric acid. Tryptamine (D4) as ISTD for tryptamine. Hippurate (D2) as ISTD for hippurate.

### Analysis of metabolites in human plasma (corresponding to Fig. 1d, Supplementary Table 5)

De-identified human plasma samples were obtained from the Stanford Blood Center under their IRB-approved research protocol. Thawed samples were mixed by gentle inversion and aliquots (50 μL) were transferred to a 96-well V-bottom plate containing internal standard mixture (tryptamine (D4), indoleacetic acid (D5), phenylacetic acid (D5), phenylpropionic acid (D9), phenylalanine (D5), hippuric acid (D2), acetic acid (^13^C_2_), isovaleric acid (D2), 40 μM each, 5 μL) and acetonitrile: methanol 3:1 mixture (150 μL). The solution was mixed five times by pipetting, then the plate was centrifuged at 5,000*g* for 15 min at 4 °C. Supernatant was collected for the following sample preparations, either derivatization or direct dilution, before being subjected to LC-MS analysis. Derivatization was performed as the protocol described above with the following minor adjustment: after reaction completion, sixteen microliters of the reaction mixture were quenched with 184 μL 0.04% formic acid in 10% acetonitrile/water. Direct dilution was performed as described above, but with the following minor adjustment: seventy microliters of supernatant were directly transferred to a 96-well V-bottom plates and diluted with LC-MS grade water (70 μL) for LC-MS analysis.

Derivatized acetate, propionate, butyrate, isobutyrate, 2-methylbutyrate, isovalerate, isocaproate, 5-aminovalerate, indoleacetate, indolepropionate, phenylacetate, phenylpropionate, 3-(4-hydroxyphenyl)acetate, 3-(4- hydroxyphenyl)propionate, hippurate, phenylacetylglycine, phenylpropionylglycine, 4- hydroxyhippurate, (3-(4-hydroxyphenyl)propanoyl)glycine, and 4- hydroxyphenylacetylglycine were analyzed with the same acquisition method as described above for mouse plasma samples, except that 20 μL of samples were injected.

### Stable isotope amino acid conversion in culture and in cell suspensions (corresponding to Fig. 2b,d, Extended data Fig. 4 and 5)

To monitor consumption of specific amino acids during growth, we prepared SACC medium containing all 20 proteinogenic amino acids and glucose (10 mM), where individual amino acids (Val, Ile, Leu, Pro, Phe, Tyr, Trp) were substituted their deuterium labeled isotopologues (Val (D8), Ile (D10), Leu (D10), Pro (D7), Phe (D8), Tyr (D4), Trp (D5)). Wild-type *C. sporogenes* were streaked from an anaerobic glycerol stock onto RCM agar plates, and incubated for ∼20 h at 37 °C. Individual well-isolated colonies were inoculated into 4 mL of SACC containing 20 AA (1 mM each) and glucose (10 mM) and cells were cultured for 20 h at 37 °C. Cells were diluted to ∼0.01 OD_600nm_ into fresh SACC with 19 unlabeled amino acid plus one stable isotope labeled amino acid (1 mM each) and glucose (10 mM) and cultured for 24 h at 37 °C. At 0 h, 6 h, and 24 h, 200 μL aliquots of culture was transferred to pre-chilled tubes and was centrifuged at ∼13,000*g* speed for 1 min at 4 °C. Supernatant was transferred to a new tube and stored at −80 °C until LC-MS measurement.

*C. sporogenes* requires 10 essential amino acids to support its growth, complicating analysis of its metabolism. To avoid interference from other amino acids, we used whole cell suspensions to measure single or paired amino acid conversions. Individual well-isolated colonies of *C. sporogenes* were inoculated into 4 mL of SACC broth plus 1 mM *trans*-4-hydroxy-L-proline and were cultured for 14 h at 37 °C. Overnight cultures were diluted to ∼0.01 OD_600nm_ into the same medium and incubated for 4 h at 37 °C until 0.15 ∼0.2 OD_600nm_. Cells were spin down anaerobically at 6,500*g* for 8 min at room temperature. Cell pellet was washed by SACC-basal solution (no amino acid nor carbon source) and resuspended in the same solution. Cell suspension was mixed with designed amino acid(s) (final 5 mM amino acid and OD_600nm_ =1 of cells if not specified) and incubate at 37 °C anaerobic incubator. For Val and Pro pair, 5 mM Val and 10 mM Pro were used to match stoichiometric requirements^15^. At T=0, 30 min, 60 min, and 180 min, assay aliquots (200 μL) were transferred to pre-chilled Eppendorf tubes. After centrifugation at ∼13,000*g* for 1 min at 4 °C, supernatant (120 μL) was transferred to a new tube and stored at −80 °C until LC-MS measurements.

#### Stable isotope-labeled amino acid and metabolite analysis by LC-MS for stable isotope amino acid conversion in growth culture and in whole cell suspension (Val, Pro)

Culture supernatants (10 μL) were mixed with internal standards (phenylethylamine (D5), phenylpropionic acid (D9), phenylalanine (D5), alanine (D4) and isovaleric acid (D2), 5 mM each, 10 μL) and diluted with LC-MS grade water (30 μL) in 96-well V- bottom plates (USA scientific cat. # 1833-9600). Then 150 μL of acetonitrile/methanol (3:1 ratio) was added. The solution was mixed five times by pipetting, then the plate was centrifuged at 5,000*g* for 10 min at 4 °C. Supernatant was collected for the following sample preparations, either derivatization or direct dilution, before subjecting to LC-MS analysis.

Derivatization for short chain fatty acid was performed as the protocol described above with slight modification: after derivatization, 2 μL of the reaction mixture were then quenched in a new 96-well V-bottom plate containing 0.02% formic acid in 10% acetonitrile/water (98 μL) and subjected to LC-MS analysis.

Direct dilution method: five microliters of supernatant were directly transferred to a 96-well V-bottom plates and diluted with LC-MS grade water (95 μL) for LC-MS analysis.

The samples were analyzed using an Agilent 6545XT Q-TOF mass spectrometer equipped with a dual jet stream electrospray ionization source operating under extended dynamic range (EDR 1700 m/z). Reverse phase separation was performed on an Agilent 1290 Infinity II LC system equipped with Waters BEH C18 1.7-μm particle size columns (2.1 × 100 mm) for all reversed phase (RP) separation.

Derivatized short chain fatty acids (acetate (D2 and D3), propionate (D3 and D5), isobutyrate (D7), butyrate (D7), 2-methyl-butyrate (D9), isovalerate (D9), isocaproate (D9 and D10)) were analyzed under negative ionization mode. Eluent A consisted of 0.1% formic acid (v/v) in water and eluent B consisted of 0.1% formic acid (v/v) in 100% acetonitrile. Samples (2 μL) were injected via refrigerated autosampler into mobile phase and separation was achieved at 40 °C using dual binary pumps with alternating column regeneration. The analytical pump applied the following gradient at a flow rate of 0.4 mL min^−1^: 0−3 min, 15% B, 3−10 min, 15% to 40% B, 10−10.1 min, 40% to 15% B, 10.1 min−11 min, 15% B. The regeneration pump applied the following gradient at a flow rate of 0.2 mL min^−1^: 0−7 min, 98% B, 7−8 min, 98% to 15% B, 8−11 min, 15% B. Electrospray ionization parameters included a gas temperature of 300 °C, drying gas flow of 6 L min^-1^, nebulizer pressure of 30 psi, sheath gas temperature of 300 °C and flow of 11 L min^-1^, capillary voltage of 4000 V, and fragmentor voltage of 140 V.

Underivatized arginine (D7), serine (D3), *trans*-4-hydroxyproline (D3 and D6), proline (D7), 5-aminovalerate (D2 and D6), valine (D8), isoleucine (D10), tyrosine (D4), leucine (D10), phenylalanine (D8), tryptamine (D5), and tryptophan (D5) were analyzed under positive ionization mode. Eluent A consisted of 0.2% formic acid (v/v) in water and eluent B consisted of 0.2% formic acid (v/v) in 100% methanol. Samples (0.5 μL) were injected via refrigerated autosampler into mobile phase and separation at 25 °C was achieved at a flow rate of 0.2 mL min^−1^ using the following gradient: 0−3 min, 1% B, 3−5 min, 1% to 98% B, 5−7 min, 98% B, 7−7.01 min, 98% to 1% B, 7.01−9.7 min, 1% Electrospray ionization parameters included a gas temperature of 300 °C, drying gas flow of 6 L min^-1^, nebulizer pressure of 30 psi, sheath gas temperature of 275 °C and flow of 11 L min^-1^, capillary voltage of 4000 V, and fragmentor voltage of 140 V.

Underivatized aromatic metabolites (4-OH-phenylpropionate (D4), 4-OH-phenylacetate (D4), indolepropionate (D5), indoleacetate (D5), phenylpropionate (D6), phenylacetate (D7)) were analyzed under negative ionization mode. Eluent A consisted of 0.1% acetic acid (v/v) in water and eluent B consisted of 0.1% acetic acid (v/v) in 100% acetonitrile. Samples (0.5 μL) were injected via refrigerated autosampler into mobile phase and separation at 40 °C was achieved at a flow rate of 0.3 mL min^−1^ using the following gradient: 0−4 min, 10% to 60% B, 4−4.5 min, 60% to 98% B, 4.5−6 min, 98% B, 6−6.01 min, 98% to 10% B, 6.01−9 min, 10% B. Electrospray ionization parameters included a gas temperature of 300 °C, drying gas flow of 6 L min^-1^, nebulizer pressure of 30 psi, sheath gas temperature of 275 °C and flow of 11 L min^-1^, capillary voltage of 4000 V, and fragmentor voltage of 140 V.

MS1 spectra were collected in centroid mode, and peak assignments in samples were made based on comparisons of retention times and accurate masses from unlabeled standards. Amino acids and metabolites were quantified by isotope-dilution mass spectrometry using corresponding ISTD. Selection of internal standards for LC-MS quantitation are as follows: phenylalanine (D5) as ISTD for arginine (D7), valine (D8), isoleucine (D10), tyrosine (D4), leucine (D10), phenylalanine (D8), tryptamine (D5), and tryptophan (D5). Alanine (D4) as ISTD for 5-aminovalerate (D2 and D6), *trans*-4-hydroxyproline (D3 and D6), proline (D7), and serine (D3). Phenylethylamine (D5) as ISTD for tryptamine (D5). Phenylpropionic acid (D9) as ISTD for 4-OH- phenylpropionate (D4), 4-OH-phenylacetate (D4), indolepropionate (D5), indoleacetate (D5), phenylpropionate (D6), and phenylacetate (D7). Isovaleric acid (D2) as ISTD for short chain fatty acids.

#### Analysis of stable isotope-labeled amino acids and metabolites in whole cells suspension (Phe, Trp, Tyr)

Culture supernatants (10 μL) were prepared using the above method with slight changes. 1) The concentration of internal standards changed as follows: phenylethylamine (D5), phenylpropionic acid (D9), phenylalanine (D5), alanine (D4), 0.2 mM each, and isovaleric acid (D2), 1 mM). 2) Aromatic amino acid metabolites (4-OH-phenylpropionate (D4), 4-OH-phenylacetate (D4), indolepropionate (D5), indoleacetate (D5), phenylpropionate (D6), and phenylacetate (D7)) were derivatized using the same method and analyzed together with short chain fatty acids. No separate analysis of underivatized metabolites under negative ionization mode was performed. Eluent A consisted of 0.1% acetic acid (v/v) in water and eluent B consisted of 0.1% acetic acid (v/v) in 100% acetonitrile. Samples (1 μL) were injected via refrigerated autosampler into mobile phase and separation was achieved with Waters BEH C18 1.7-μm particle size C18 columns (2.1 × 100 mm) at 40 °C using dual binary pumps with alternating column regeneration. The analytical pump applied the following gradient at a flow rate of 0.4 mL min^−1^: 0−4 min, 3% to 50% B, 4−4.1 min, 50% to 3% B, 4.1−5 min, 3% B. The regeneration pump applied the following gradient at a flow rate of 0.4 mL min^−1^: 0−0.1 min, 3% to 98% B, 0.1−2 min, 98% B, 2−2.2 min, 98% to 3% B, 2.2−5 min, 3% B. Electrospray ionization parameters included a gas temperature of 250 °C, drying gas flow of 6 L min^-1^, nebulizer pressure of 30 psi, sheath gas temperature of 250 °C and flow of 11 L min^-1^, capillary voltage of 4000 V, and fragmentor voltage of 70 V.

Selection of internal standards for LC-MS quantitation was as follows: phenylalanine (D5) as ISTD for arginine (D7), valine (D8), isoleucine (D10), tyrosine (D4), leucine (D10), methionine (D8), phenylalanine (D8), tryptamine (D5), and tryptophan (D5). Alanine (D4) as ISTD for 5-aminovalerate (D2 and D6), *trans*-4- hydroxyproline (D3 and D6), proline (D7), and serine (D3). Phenylethylamine (D5) as ISTD for tryptamine (D5). Phenylpropionic acid (D9) as ISTD for 4-OH-phenylpropionate (D4), 4-OH-phenylacetate (D4), indolepropionate (D5), indoleacetate (D5), phenylpropionate (D6), and phenylacetate (D7). Isovaleric acid (D2) as ISTD for short chain fatty acids.

### ATP measurements in whole cells and detection of metabolites (corresponding to Fig. 3d, Extended data Fig. 6 and 7)

Wild type *C. sporogenes* cells were streaked from an anaerobic glycerol stock onto a TYG (3% w/v tryptone, 2% w/v yeast extract, 0.1% w/v sodium thioglycolate) agar plate and incubated at 37 °C in an anaerobic chamber from Coy Laboratories under an atmosphere of 5% H_2_, 10% CO_2_ and 85% N_2_. After ∼24 h, a well isolated colony was inoculated into 5 mL of TYG broth and cultured to stationary phase (∼24 h). Cells were diluted into 20 mL of TYG broth (1000-fold dilution) and grown to late-log phase of growth (∼16 h). The cells were then harvested by centrifugation (5,000*g*, 10 min, 4 °C) and washed twice with 20 mL volumes of pre-reduced phosphate assay buffer (40 mM potassium phosphate, 10 mM magnesium sulfate, pH 7.0). All centrifugation steps were performed outside of the anaerobic chamber, and so to ensure anaerobiosis, 50 mL falcon tubes were tightly capped and care was taken to limit the total amount of time outside of the chamber. After the final wash step, the cell pellet was re-suspended in 1 ml of phosphate assay buffer and incubated at room temperature for 45 min to achieve a resting cell suspension. During this time, 100 μL of cells were seeded into rows of a 96-well microtiter plate (12 wells per condition being tested). Two hundred microliters of pre-reduced 2 mM substrate in phosphate assay buffer, or buffer alone was dispensed into rows of a separate 96-well microplate. At time zero, 100 μL of substrate or buffer was added to the cells and mixed gently by pipetting. At - 5 min, - 1 min, +30 s, +1 min, +2 min, +5 min, +10 min, +20 min, +30 min, +45 min, +60 min, and +90 min, 10 μL of cells were taken and immediately mixed with 90 μL of DMSO to quench the reaction and liberate cellular ATP. The −5 and −1 minute time-points were taken prior to addition of buffer or substrate, so only 5 μL of cell suspension was harvested, and 5 μL of either buffer or substrate was added to the cell-DMSO mixtures to bring the total volume to 100 μL. At the end of the assay, a 50 μL aliquot of cell suspension was taken for subsequent total protein determination. The ATP content from 10 μL aliquots of lysed cells was then detected using a luminescence-based ATP determination kit from Life Technologies and ATP levels were calculated by extrapolation from a calibration curve constructed with known concentrations of ATP. For protein determination, cellular protein was solubilized in NaOH by combining equal volumes of cells and 0.4 M NaOH (50 μL, each) and incubating at 99 °C for 10 minutes. The protein content was then determined using the DC Protein Assay Kit II from BioRad using bovine serum albumin as a standard. ATP levels were expressed as nmol of ATP per mg of total protein.

To test the effect of 3,3’,4’,5-tetrachlorosalicylanilide (TCS) on ATP formation, washed cells first were rested at room temperature anaerobically for 30 min. Then TCS (diluted from a 100 X stock in ethanol to achieve the desired concentration) was added and cells were incubated at room temperature for an additional 30 min. Assays were then performed as described above, and ten microliters of DMSO quenched samples were used immediately for ATP measurement, with the remaining 90 μL being stored at −80 °C until LC-MS measurements.

Before LC-MS measurement, DMSO quenched samples (90 μL) were thawed at room temperature and then centrifuged at 5,000*g* for 15 min at room temperature.

For quantitation of phenylalanine, phenylethylamine, phenyllactate, phenylpyruvate, trans-cinnamate, PPA, PPG, and PAA, supernatant (60 μL) was mixed with internal standard solution (PPA (D9), phenylalanine (D5), phenylethylamine (D5), PAA (D5), 100 μM each, 6 μL). Twenty-five microliters of the above mixture were diluted with 75 μL LC-MS grade water. Eluent A consisted of 0.1% acetic acid (v/v) in water and eluent B consisted of 0.1% acetic acid (v/v) in 100% acetonitrile. Samples (1 μL) were injected via refrigerated autosampler into mobile phase and separation with a BEH C18 1.7-μm particle size C18 column (2.1 × 100 mm) at 40 °C was achieved at a flow rate of 0.4 mL min^−1^ with the following gradient: 0−1 min, 3% B, 1−4 min, 3% to 60% B, 4−4.1 min, 60% to 99.9% B, 4.1−7 min, 99.9% B, 7−7.1 min, 99.9% to 3% B, 7.1−9 min, 3% B. Two-time segments were applied for mass spectra acquisition. Segment 1 from 0.5 to 2.65 min was set under positive ionization mode for analysis of phenylalanine and phenylethylamine. Segment 2 from 2.65 to 5 min was set under negative ionization mode for analysis of other metabolites. Electrospray ionization parameters included a gas temperature of 250 °C, drying gas flow of 6 L min^-1^, nebulizer pressure of 30 psi, sheath gas temperature of 250 °C and flow of 11 L min^-1^, capillary voltage of 4000 V, and fragmentor voltage of 70 V.

For quantitation of proline and 5-aminovalerate, supernatant (60 μL) was mixed with internal standard (phenylalanine (D5), 100 μM, 6 μL). Twenty-five microliters were diluted with 50% acetonitrile containing 0.1% acetic acid (75 μL) for quantitation of proline and 5-aminovalerate. The analytes were measured with hydrophilic interaction liquid chromatography (HILIC) using a Waters BEH amide 1.7-μm particle size column (2.1 × 100 mm). Eluent A consisted of 5 mM ammonium acetate, 0.1% acetic acid (v/v) in water, and eluent B consisted of 5 mM ammonium acetate, 0.1% acetic acid (v/v) in 90% acetonitrile. Samples (2 μL) were injected via refrigerated autosampler into mobile phase and HILIC separation at 50 °C was achieved at a flow rate of 0.4 mL min^−1^ with the following gradient: 0−1 min, 80% B, 1−2.5 min, 80% to 40% B, 2.5−3.5 min, 40% B, 3.5−4 min, 40% to 80% B, 4−7 min, 80% B. Mass spectra were acquired under negative ionization mode with the following parameters: gas temperature of 250 °C, drying gas flow of 6 L min^-1^, nebulizer pressure of 30 psi, sheath gas temperature of 250 °C and flow of 11 L min^-1^, capillary voltage of 4000 V, and fragmentor voltage of 70 V.

MS1 spectra were collected in centroid mode, and peak assignments in samples were made based on comparisons of retention times and accurate masses from standards. Amino acids and metabolites were quantified by isotope-dilution mass spectrometry using corresponding ISTD. Selection of internal standards for LC-MS quantitation are as follows: phenylalanine (D5) as ISTD for arginine (D7), valine (D8), isoleucine (D10), tyrosine (D4), leucine (D10), phenylalanine (D8), tryptamine (D5), and tryptophan (D5).

### Colonization of gnotobiotic mice with wild-type or rnfB mutant C. sporogenes and analysis of indolepropionate in cecal contents and serum (corresponding to Fig. 4e)

Mouse experiments were performed on gnotobiotic Swiss Webster germ-free mice (male, 6–10 weeks of age, n = 5 per group) originally obtained from Taconic Biosciences maintained in aseptic isolators. Animal experiments were performed following a protocol approved by the Stanford University Administrative Panel on Laboratory Animal Care. Separate groups of mice were colonized with wild-type or *rnfB* mutant *C. sporogenes* by oral gavage (200 μL, ∼1 × 10^7^ CFU) and housed in separate flexible film isolators. After 2 weeks, mice were euthanized and blood was collected by cardiac puncture, placed in serum separator tubes (BD cat. # 365967), mixed briefly, incubated at room temperature for ∼30 min, then serum was obtained by centrifuging tubes at 10,000*g* for 10 min at room temperature. Cecal contents were also collected and along with serum aliquots were stored at −80 °C until analysis. Indolepropionate was then quantified by LC-MS as described in the section above for metabolite analysis in body fluids of gnotobiotic mice.

### Competition experiments for wild-type and rnfB mutant C. sporogenes in gnotobiotic mice

Mouse experiments were performed on gnotobiotic Swiss Webster germ-free mice (male, 6–10 weeks of age, n = 5 per group) originally obtained from Taconic Biosciences maintained in aseptic isolators. Animal experiments were performed following a protocol approved by the Stanford University Administrative Panel on Laboratory Animal Care. For the mono-colonization experiments, mice were colonized with a 1:1 mixture of wild-type and *rnfB* mutant *C. sporogenes* by oral gavage (200 μL, ∼1 × 10^7^ CFU) and were maintained on standard chow (LabDiet 5k67). For the defined community, mice were first colonized with a mixture consisting of equal volumes from cultures of *Bacteroides thetaiotaomicron* (∼16 h overnight culture), *Bacteroides vulgatus* (∼ 16 h overnight culture), *Edwardsiella tarda* (∼ 16 h overnight culture), *Eubacterium rectale* (∼24 h overnight culture), *Clostridium scindens* (∼30 h culture), and *Bifidobacterium breve* (∼24 h overnight culture). After one week, mice were colonized with a 1:1 mixture of wild--type or *rnfB* mutant *C. sporogenes* (200 μL, ∼1 × 10^7^ CFU) and were maintained on standard chow (LabDiet 5k67). Fresh fecal pellets were collected and stored at −80 °C until DNA extraction was performed.

#### DNA isolation and Q-PCR

Relative abundances of the wild-type and *rnfB C. sporogenes* strains over time following colonization was determined by Q-PCR (*corresponding to* ***Fig. 4f,g***). Approximately 10-20 mg of feces was added to individual tubes of a DNeasy PowerSoil DNA Isolation Kit (formerly MO BIO) and DNA was extracted using a Retsch MM400 ball mill according to manufacturer’s guidelines. DNA quantitation was performed with an Invitrogen Quant-iT broad range dsDNA assay kit. Two primer sets were used, one designed to amplify total *C. sporogenes*, and another specifically targeted to the erythromycin resistance cassette of the *rnfB* mutant (**Supplementary Table 8**). To amplify genomic DNA (gDNA), we used the Brilliant III SYBR kit from Agilent Technologies. Conditions were as follows: 96-well Q-PCR plates, 20 μL reactions, primers at 500 nM each, reference dye included, 5 ng DNA per reaction. Amplification was performed in duplicate using a Stratagene Mx3000P with PCR conditions supplied in the Brilliant III SYBR kit. Initial validation plots were performed using eight serial 4-fold dilutions of gDNA from either WT or *rnfB* mutant *C. sporogenes* spanning 50 ng to 3 pg per reaction. The cycle threshold vs. log(input DNA) was plotted and a linear curve fit was applied. Both primers displayed excellent linearity across this range of DNA concentrations (R^2^>0.98). For fecal DNA samples, the cycle number at the threshold crossing point was used to calculate the amount of total *C. sporogenes* DNA or *rnfB* mutant DNA using the standard curves. The competitive index was calculated according to equation 1 below:

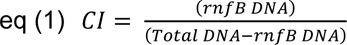

Relative compositions of community members (*corresponding to **Extended data*** ***Fig. 10***) were measured by Q-PCR using PowerUp SYBR Green Master Mix (Thermo Fisher, A25742), 400 nM primers, 5∼15 ng DNA per reaction in 384-well Q-PCR plates (Applied Biosystems™ Cat. # 4483285) with total 12 μL reaction. Amplification was performed in triplicate using QuantStudio™ 5 according to SYBR Green user manual following by melting curve. Primer validation were performed using six serial 10-fold dilutions of gDNA from each strain, spanning 24 ng to 0.24 pg per reaction. Primer amplification efficiency were between 92∼100%. For fecal DNA samples, the amount of each strains was calculated using the standard curves. The relative composition was calculated according to equation 2 below:

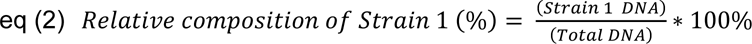

### Bioinformatics analysis (corresponding to Extended data Fig. 11, 12 and Supplementary Table 6)

#### Analysis of Rnf clusters in reference genomes from the human microbiome project

Reference genomes (GenBank file format) from the NIH Human Microbiome Project (BioProject ID 43021) were downloaded on June 19^th^, 2019 which included 2,284 genome sequences. We then used the software tool MultiGeneBlast^30^ to search these reference genomes for homologs of the *C. sporogenes* Rnf gene cluster (*rnfCDGEAB*, CLOSPO_00568 to CLOSPO_00573) with default settings. The results were manually curated to identify organisms harboring complete gene clusters. Coding sequences for the 16S rRNA genes were retrieved for these organisms from the Joint Genome Institute Integrated Microbial Genomes server. Sequences were uploaded into Geneious Prime (v2019.1.3) and were aligned using MUSCLE with default settings. Alignments were trimmed to remove incomplete sequences (<500 bp) and gaps, then were re-aligned and a phylogenetic tree was constructed using the neighbor joining method. Representative gene clusters were exported from MultiGeneBlast and visualized in Adobe Illustrator (v25.1).

#### Analysis of electron transfer proteins in GenBank

To identify homologs of the electron transfer proteins analyzed in this study, we performed BLASTp searches of the GenBank database. Query sequences included the butyryl-CoA dehydrogenase (Bcd) from *C. kluyveri* DSM 555 (GenBank acc. # EDK32509.1), the (aryl)acryloyl-CoA dehydrogenase (AcdA) from *C. sporogenes* ATCC 15579 (GenBank acc. # EDU39257.1), the isocaprenoyl-CoA dehydrogenase (AcdB) from *C. sporogenes* ATCC 15579 (GenBank acc. # EDU36591.1), the proline reductase subunit A (PrdA) from *C. sticklandii* DSM 519 (GenBank acc. # CBH22353.1), and the RnfB protein from *C. sporogenes* ATCC 15579 (GenBank acc. # EDU37753.1). Our initial analysis of gene cluster neighborhoods for these homologs using MultiGeneBlast suggested a cutoff of 65% amino acid identify was necessary to identify likely isofunctional homologs for Bcd, AcdA, AcdB, and PrdA, whereas a less stringent cutoff of 50% amino acid identity was suitable for RnfB. These query sequences were used in BLASTp searches of the GenBank database (December 30^th^, 2020) and homologs with less than 80% coverage over the length of the proteins were discarded.

#### Analysis of abundance and prevalence of electron transfer proteins in metagenomics datasets

To assess the abundance and prevalence of electron transfer proteins (Bcd, AcdA, AcdB, PrdA, and RnfB) in human fecal metagenomic datasets, we employed the web-based tool, MetaQuery^31^. Query sequences included the butyryl-CoA dehydrogenase (Bcd) from *C. kluyveri* DSM 555 (GenBank acc. # EDK32509.1), the (aryl)acryloyl-CoA dehydrogenase (AcdA) from *C. sporogenes* ATCC 15579 (GenBank acc. # EDU39257.1), the isocaprenoyl-CoA dehydrogenase (AcdB) from *C. sporogenes* ATCC 15579 (GenBank acc. # EDU36591.1), the proline reductase subunit A (PrdA) from *C. sticklandii* DSM 519 (GenBank acc. # CBH22353.1), and the RnfB protein from *sporogenes* ATCC 15579 (GenBank acc. # EDU37753.1). Owing to limitations in threshold selections implemented in MetaQuery, we used a slightly lower threshold for amino acid percent identity of 60% for Bcd, AcdA, AcdB, and PrdA to identify homologs in metagenomics datasets.

## Data availability

The authors declare that the data supporting the findings of this study are available within the paper and its Supplementary Information.

## Acknowledgements

We are grateful to Michael Fischbach, Christopher Walsh, Audrey Southwick and Curt Fischer (Stanford) and Isaac Cann (Illinois) for valuable discussions. We thank Manhong Wu for assistance with amino acid analysis by LC-MS, Michelle St. Onge with help constructing mutants, and Chun-Jun Guo and Ricardo De La Pena for assistance with GC-MS analysis of short chain fatty acids. This work was funded in part by National Institutes of Health grants DK110335 (D.D.), DK101674 (J.L.S.), DK085025 (J.L.S.) and the Chan Zuckerberg Biohub (J.L.S.).

## References

1. Nisman, B. The Stickland reaction. Bacteriol Rev 18, 16–42 (1954).

2. Smith, E. A. & Macfarlane, G. T. Dissimilatory amino acid metabolism in human colonic bacteria. Anaerobe 3, 327–337 (1997).

3. Russell, W. R. et al. Major phenylpropanoid-derived metabolites in the human gut can arise from microbial fermentation of protein. Mol Nutr Food Res 57, 523–535 (2013).

4. Van Treuren, W. & Dodd, D. Microbial contribution to the human metabolome: Implications for health and disease. Annu Rev Pathol (2019).

5. Liu, Y., Hou, Y., Wang, G., Zheng, X. & Hao, H. Gut microbial metabolites of aromatic amino acids as signals in host-microbe interplay. Trends Endocrinol Metab 31, 818–834, doi:10.1016/j.tem.2020.02.012 (2020).

6. Dodd, D. et al. A gut bacterial pathway metabolizes aromatic amino acids into nine circulating metabolites. Nature 551, 648–652 (2017).

7. Venkatesh, M. et al. Symbiotic bacterial metabolites regulate gastrointestinal barrier function via the xenobiotic sensor PXR and Toll-like receptor 4. Immunity 41, 296–310, doi:10.1016/j.immuni.2014.06.014 (2014).

8. Roager, H. M. & Licht, T. R. Microbial tryptophan catabolites in health and disease. Nat Commun 9, 3294, doi:10.1038/s41467-018-05470-4 (2018).

9. Steed, A. L. et al. The microbial metabolite desaminotyrosine protects from influenza through type I interferon. Science 357, 498–502, doi:10.1126/science.aam5336 (2017).

10. Allison, M. J., Bryant, M. P. & Doetsch, R. N. Volatile fatty acid growth factor for cellulolytic cocci of bovine rumen. Science 128, 474–475 (1958).

11. Stack, R. J., Hungate, R. E. & Opsahl, W. P. Phenylacetic acid stimulation of cellulose digestion by *Ruminococcus albus* 8. Appl Environ Microbiol 46, 539–544 (1983).

12. Hungate, R. E. & Stack, R. J. Phenylpropanoic acid: Growth factor for *Ruminococcus albus*. Appl Environ Microbiol 44, 79–83 (1982).

13. Stickland, L. H. Studies in the metabolism of the strict anaerobes (genus *Clostridium*): The chemical reactions by which *Cl. sporogenes* obtains its energy. Biochem J 28, 1746–1759 (1934).

14. Barker, H. A. Amino acid degradation by anaerobic bacteria. Annu Rev Biochem 50, 23–40 (1981).

15. Lovitt, R. W., Kell, D. B. & Morris, J. G. The physiology of *Clostridium sporogenes* NCIB 8053 growing in defined media. J Appl Bacteriol 62, 81–92 (1987).

16. Nemet, I. et al. A cardiovascular disease-linked gut microbial metabolite acts via adrenergic receptors. Cell 180, 862–877 e822, doi:10.1016/j.cell.2020.02.016 (2020).

17. Wildenauer, F. X. & Winter, J. Fermentation of isoleucine and arginine by pure and syntrophic cultures of *Clostridium sporogenes*. FEMS Microbiol Lett 38, 373–379 (1986).

18. Macy, J., Probst, I. & Gottschalk, G. Evidence for cytochrome involvement in fumarate reduction and adenosine 5’-triphosphate synthesis by *Bacteroides fragilis* grown in the presence of hemin. J Bacteriol 123, 436–442, doi:10.1128/JB.123.2.436-442.1975 (1975).

19. Hess, V., Gonzalez, J. M., Parthasarathy, A., Buckel, W. & Muller, V. Caffeate respiration in the acetogenic bacterium *Acetobacterium woodii*: a coenzyme A loop saves energy for caffeate activation. Appl Environ Microbiol 79, 1942–1947 (2013).

20. Bader, J. & Simon, H. ATP formation is coupled to the hydrogenation of 2-enoates in *Clostridium sporogenes*. FEMS Microbiol Lett 20, 171–175 (1983).

21. Tremblay, P. L., Zhang, T., Dar, S. A., Leang, C. & Lovley, D. R. The Rnf complex of *Clostridium ljungdahlii* is a proton-translocating ferredoxin:NAD+ oxidoreductase essential for autotrophic growth. mBio 4, e00406–00412, doi:10.1128/mBio.00406-12 (2012).

22. Lovitt, R. W., Kell, D. B. & Morris, J. G. Proline reduction by *Clostridium sporogenes* is coupled to vectorial proton ejection. FEMS Microbiol Lett 36, 269–273 (1986).

23. Seedorf, H. et al. The genome of *Clostridium kluyveri*, a strict anaerobe with unique metabolic features. Proc Natl Acad Sci USA 105, 2128–2133 (2008).

24. Li, F. et al. Coupled ferredoxin and crotonyl coenzyme A (CoA) reduction with NADH catalyzed by the butyryl-CoA dehydrogenase/Etf complex from *Clostridium kluyveri*. J Bacteriol 190, 843–850 (2008).

25. Buckel, W. & Thauer, R. K. Flavin-based electron bifurcation, a new mechanism of biological energy coupling. Chem Rev 118, 3862–3886 (2018).

26. Herrmann, G., Jayamani, E., Mai, G. & Buckel, W. Energy conservation via electron- transferring flavoprotein in anaerobic bacteria. J Bacteriol 190, 784–791 (2008).

27. Guo, C. J. et al. Depletion of microbiome-derived molecules in the host using *Clostridium* genetics. Science 366, doi:10.1126/science.aav1282 (2019).

28. Goodman, A. L. et al. Identifying genetic determinants needed to establish a human gut symbiont in its habitat. Cell Host Microbe 6, 279–289, doi:10.1016/j.chom.2009.08.003 (2009).

29. Wu, M. et al. Genetic determinants of in vivo fitness and diet responsiveness in multiple human gut *Bacteroides*. Science 350, aac5992, doi:10.1126/science.aac5992 (2015).

30. Medema, M. H., Takano, E. & Breitling, R. Detecting sequence homology at the gene cluster level with MultiGeneBlast. Mol Biol Evol 30, 1218–1223, doi:10.1093/molbev/mst025 (2013).

31. Nayfach, S., Fischbach, M. A. & Pollard, K. S. MetaQuery: a web server for rapid annotation and quantitative analysis of specific genes in the human gut microbiome. Bioinformatics 31, 3368–3370, doi:10.1093/bioinformatics/btv382 (2015).

