## Supplementary Information for "Electron transfer proteins in gut bacteria yield metabolites that circulate in the host"

### Supplementary Text

#### *Growth based assay to study Stickland metabolism in C. sporogenes*

*Clostridium sporogenes* is a proteolytic bacterium that obtains its energy by coupling the oxidation of one amino acid with reduction of another (the Stickland reaction). Several amino acid pairs are known to stimulate its growth including Arg/Ile[1], Ser/Pro, Val/Pro, Leu/Pro, and Ile/Pro[2]. However, a comprehensive analysis of the growth promoting Stickland pairs has not been performed. To assess the degree to which amino acid pairs are used to promote growth, we performed a high throughput growth-based screen. We used a defined medium which contains ten amino acids required for growth (Gly, Val, Leu, Ile, Met, His, Arg, Phe, Tyr, Trp; 1 mM each), trace elements, minerals, and vitamins. To this medium, we added higher concentrations (25 mM each) of substrates (amino acids or glucose) either individually or in pairwise combinations. Thus, this system is designed to detect conditions in which pairs of amino acids enable cells to reach higher optical densities compared to basal medium alone or individual substrates. In initial experiments, we failed to observe growth stimulation by the addition of valine and proline. As previously reported[2], we found that this was due to a requirement for acetate to be supplemented in the medium, likely to satisfy anabolic needs. Thus, we performed growth-based screens under conditions where acetate was either added (40 mM) or left out from the basal medium.

Under these conditions, a finite number of substrate combinations enabled large increases in growth (**Extended data Fig. 2a**). Growth from glucose was most robust when combined with reductive pathway substrates such as proline or pathways that converge on proline (e.g., trans-4-hydroxyproline, arginine, citrulline, ornithine).

Previous studies have indicated that *C. sporogenes* metabolizes glucose via the Embden-Meyerhof-Parnas pathway with acetate and ethanol being major fermentation products[3]. Proline supplementation diminishes ethanol production, the reducing equivalents most likely being disposed of via 5-aminovalerate production. This 'mixed Stickland' reaction improves maximal growth rate and molar growth yield on glucose, suggesting that proline reduction is involved in energy conservation. In this context, our observation that additional substrates *trans*-4-hydroxyproline, arginine, citrulline, and ornithine also improve growth yields from glucose suggests a convergence of metabolism in *C. sporogenes* on the proline reductase complex. The genome sequence for *C. sporogenes* contains homologs for the pathway converting *trans*-4-hydroxyproline to proline (*trans*-4-hydroxy-L-proline dehydratase, CLOSPO_00435-CLOSPO_00436; pyrroline-5-carboxylate reductase, CLOSPO_00437), and arginine to proline via citrulline and ornithine (arginine deiminase, CLOSPO_03346 or CLOSPO_00894; Ornithine carbamoyltransferase, CLOSPO_02415; Carbamate kinase, CLOSPO_02414; Ornithine cyclodeaminase, CLOSPO_02434; Proline racemase, CLOSPO_02527). Another proteolytic gut bacterium, *Clostridioides difficile* also displays a propensity for proline reduction[4,5], suggesting that this mechanism may be common among amino acid utilizing bacteria in the gut.

Cells incubated with serine showed a very similar pattern to glucose with respect to the interaction with other amino acids (**Extended data Fig. 2a**). Serine is deaminated in a single reaction to form pyruvate, a central metabolic intermediate that can be oxidatively decarboxylated, yielding energy via substrate level phosphorylation. In this respect, serine metabolism shares similarity with glucose metabolism, both involving pyruvate as a key metabolic intermediate. The genome of *C. sporogenes* encodes a putative serine dehydratase (CLOSPO_03760-CLOSPO_03761). This enzyme is unique among ammonia-lyases in that it employs an oxygen-labile iron sulfur cluster for catalysis rather than pyridoxal phosphate[6]. In our assay, the Ser/Arg pair promoted the largest growth yield of any combination of amino acids or glucose (**Extended data Fig. 2a**). Quantitative growth curves performed using Balch-type tubes and a conventional spectrophotometer (**Extended data Fig. 2b**) revealed that the Ser/Arg combination reached a maximum OD approximately 3-fold higher than that of either of the substrates alone (**Extended data Fig. 2c**). The Ser/Arg combination did not require supplemental acetate (**Extended data Fig. 2a** vs. **Extended data Fig. 2d**), likely reflecting the capacity of serine to satisfy anabolic requirements via pyruvate. Despite the observation that most bacteria carry a copy of serine dehydratase[7], relatively little is known about its role in metabolism. In *E. coli*, serine dehydratase is thought to regulate intracellular serine levels, as mutants display aberrantly elevated serine levels, altered one-carbon metabolism and defective cell wall synthesis[8]. In asaccharolytic *Campylobacter jejuni*, serine is preferentially utilized as a growth substrate[9] and a serine dehydratase mutant is attenuated for colonization in the avian gut[10]. Our results provide additional insights into the role of serine in bacterial metabolism, suggesting that it is an important oxidative substrate for Stickland metabolism.

Acetate supplementation had a dramatic impact on certain amino acid pairs. Among the top ten Stickland pairs that were stimulated by addition of acetate, eight included branched chain amino acids (reductants) and proline or amino acids that converge on proline (oxidants) (**Extended data Fig. 2e**, red bars). Collectively, this suggests that neither branched chain amino acids nor proline and related amino acids are utilized by the cell as sources of anabolic precursors. These findings were further reinforced by our demonstration that cell suspensions incubated with stable isotope labeled proline and valine showed stoichiometric conversion to the respective metabolic products (5-aminovalerate and isobutyrate) (**Fig. 2d**).

#### *ATP formation during cinnamate reduction by C. sporogenes*

Based on earlier biochemical work, cinnamate reductase was proposed to be involved in reductive Stickland metabolism of phenylalanine and tyrosine[11]. This enzyme catalyzes the reduction of cinnamic acid and *p*-coumaric acid, predicted products of the phenyllactoyl-CoA dehydratase involved in reductive amino acid metabolism of Phe and Tyr[12]. However, cinnamate reductase is not localized to the gene cluster for reductive amino acid metabolism, and instead, a separate enzyme (acyl-CoA dehydrogenase) is located within the gene cluster that performs a similar overall reaction (**Fig. 3b**). Cinnamate reductase is located in a different chromosomal locus with the only linked gene being a *merR*-like transcriptional regulator (**Extended data Fig. 5a**). Acyl-CoA dehydrogenases are clearly distinct from cinnamate (or enoate) reductases, sharing no significant amino acid sequence homology. Whereas acyl-CoA dehydrogenases are heterotrimeric electron bifurcating flavoenzymes, enoate reductases possess an old yellow enzyme domain (FMN containing) fused with domains that carry additional flavin adenine dinucleotide and iron sulfur cluster prosthetic groups (**Extended data Fig. 5b**)[13]. Using genetics, we previously found that cinnamate reductase is involved in reduction of cinnamate and *p*-coumarate, but is not involved in aromatic amino acid metabolism[14]. Therefore, cinnamate reductase most likely functions to reduce secondary plant metabolites which include cinnamate and *p*-coumarate. Intriguingly, cinnamate reduction has previously been shown to drive ATP formation in *C. sporogenes* cells[15]. Using the ATP assay described in the methods section, we measured ATP levels after pulsing resting cell suspensions with cinnamate. After addition of cinnamate, we observed a rapid rise in ATP formation followed by a short plateau and a rapid decrease back to starting levels (**Extended data Fig. 5c**). This result, which replicates that from previous studies, provides evidence that cinnamate reductase is involved in energy conservation. The mechanism is likely distinct from that for caffeate respiration (a structurally similar plant metabolite) in *Acetobacterium woodii* which conserves energy via the electron bifurcating caffeoyl-CoA dehydrogenase[16].

### Supplementary Tables

#### Supplementary Table 1. Metabolite levels in *C. sporogenes* WT mono-colonized mouse fecal samples

| Metabolites (nmol/mg)[a] | LLOQ[b] (nmol/mg) | Germ-free control[c,d] | Before gavage[c,d] | Colonized 1 week[c,d] | Colonized 2 weeks[c,d] | Colonized 2 weeks + antibiotics[c,d] |
|---|---|---|---|---|---|---|
| 5-Aminovalerate (5-AVA) | 0.019 | n/d | n/d | 9.69 ± 0.92 | 8.44 ± 1.47 | n/d |
| Acetate | 0.30 | <LLOQ | <LLOQ | 2.89 ± 0.82 | 1.88 ± 0.36 | <LLOQ |
| Butyrate | 0.019 | <LLOQ | <LLOQ | <LLOQ | <LLOQ | <LLOQ |
| Hippurate | 0.005 | n/d | n/d | <LLOQ | <LLOQ | n/d |
| 4-Hydroxyhippurate | 0.009 | n/d | n/d | n/d | n/d | n/d |
| 4-Hydroxyphenylacetate (4-OH-PAA) | 0.005 | <LLOQ | <LLOQ | 0.008 ± 0.002 | <LLOQ | <LLOQ |
| 4-Hydroxyphenylacetylglycine (4-OH-PAG) | 0.005 | n/d | n/d | n/d | n/d | n/d |
| 4-Hydroxyphenylpropionate (4-OH-PPA) | 0.009 | n/d | n/d | 0.66 ± 0.16 | 0.38 ± 0.20 | n/d |
| 4-Hydroxyphenylpropionylglycine (4-OH-PPG) | 0.005 | n/d | n/d | n/d | n/d | n/d |
| Indoleacetate (IAA) | 0.005 | n/d | n/d | n/d | n/d | n/d |
| Indolepropionate (IPA) | 0.009 | n/d | n/d | 0.06 ± 0.01 | 0.042 ± 0.01 | n/d |
| Isobutyrate | 0.009 | <LLOQ | <LLOQ | 0.39 ± 0.08 | 0.20 ± 0.07 | <LLOQ |
| Isocaproate | 0.009 | n/d | n/d | 0.06 ± 0.03 | <LLOQ | n/d |
| Isovalerate | 0.019 | <LLOQ | <LLOQ | 0.40 ± 0.09 | 0.26 ± 0.08 | <LLOQ |
| 2-Methylbutyrate | 0.019 | n/d | n/d | 0.38 ± 0.07 | 0.20 ± 0.06 | n/d |
| Phenylacetate (PAA) | 0.005 | <LLOQ | <LLOQ | 0.01 ± 0.004 | 0.005 ± 0.002 | <LLOQ |
| Phenylacetylglycine (PAG) | 0.005 | n/d | n/d | n/d | n/d | n/d |
| Phenylpropionate (PPA) | 0.005 | n/d | n/d | 0.20 ± 0.05 | 0.12 ± 0.04 | n/d |
| Phenylpropionylglycine (PPG) | 0.002 | n/d | n/d | n/d | n/d | n/d |
| Propionate | 0.048 | <LLOQ | <LLOQ | 0.28 ± 0.07 | 0.14 ± 0.04 | <LLOQ |
| Tryptamine | 0.002 | n/d | n/d | 0.01 ± 0.005 | <LLOQ | n/d |

[a] Data reported in the table were normalized by wet weight of feces (40 ± 4 mg).

[b] LLOQ is the lower limit of quantitation for the assay.

[c] <LLOQ indicates the value is lower than lower limit of quantitation; n/d indicates the value is below assay detection limit.

[d] Data are presented as means ± standard errors from the mean. For germ-free controls, before gavage and colonized 1 week, n = 10 mice per group. For colonized 2 weeks and colonized 2 weeks + antibiotic water, n = 5 mice per group.

**Supplementary Table 2. Metabolite levels in *C. sporogenes* WT mono-colonized mouse plasma samples**

| Metabolites (μM) | LLOQ[a] (μM) | Germ-free control[b,c] | Before gavage[b,c] | Colonized 1 week[b,c] | Colonized 2 weeks[b,c] | Colonized 2 weeks + antibiotics[b,c] |
|---|---|---|---|---|---|---|
| 5-Aminovalerate (5-AVA) | 2.32 | n/d | n/d | 11.8 ± 5.14 | 11.5 ± 2.56 | n/d |
| Acetate | 148 | 358 ± 35.0 | 352 ± 59.3 | 373 ± 54.0 | 370 ± 40.6 | 483 ± 51.9 |
| Butyrate | 4.64 | <LLOQ | <LLOQ | <LLOQ | <LLOQ | <LLOQ |
| Hippurate | 4.64 | n/d | n/d | 12.4 ± 4.79 | 8.67 ± 2.40 | n/d |
| 4-Hydroxyhippurate | 2.32 | n/d | n/d | n/d | n/d | n/d |
| 4-Hydroxyphenylacetate (4-OH-PAA) | 2.32 | n/d | n/d | n/d | n/d | n/d |
| 4-Hydroxyphenylacetylglycine (4-OH-PAG) | 0.58 | n/d | n/d | n/d | n/d | n/d |
| 4-Hydroxyphenylpropionate (4-OH-PPA) | 1.16 | n/d | n/d | 13.1 ± 3.47 | 13.8 ± 0.91 | n/d |
| 4-Hydroxyphenylpropionylglycine (4-OH-PPG) | 1.16 | n/d | n/d | n/d | n/d | n/d |
| Indoleacetate (IAA) | 1.16 | n/d | n/d | <LLOQ | <LLOQ | n/d |
| Indolepropionate (IPA) | 1.16 | n/d | n/d | 91.4 ± 15.0 | 106 ± 19.5 | n/d |
| Isobutyrate | 2.32 | <LLOQ | <LLOQ | <LLOQ | <LLOQ | <LLOQ |
| Isocaproate | 2.32 | n/d | n/d | n/d | n/d | n/d |
| Isovalerate | 2.32 | <LLOQ | <LLOQ | n/d | n/d | n/d |
| 2-Methylbutyrate | 4.64 | n/d | n/d | n/d | n/d | n/d |
| Phenylacetate (PAA) | 1.16 | n/d | n/d | n/d | n/d | n/d |
| Phenylacetylglycine (PAG) | 1.16 | <LLOQ | <LLOQ | <LLOQ | <LLOQ | <LLOQ |
| Phenylpropionate (PPA) | 1.16 | n/d | n/d | 2.69 ± 0.60 | 3.04 ± 0.38 | n/d |
| Phenylpropionylglycine (PPG) | 1.16 | n/d | n/d | n/d | n/d | n/d |
| Propionate | 9.28 | <LLOQ | 9.71 ± 3.39 | 10.6 ± 3.42 | <LLOQ | <LLOQ |
| Tryptamine | 0.58 | n/d | n/d | n/d | n/d | n/d |

[a] LLOQ is the lower limit of quantitation for the assay.

[b] <LLOQ indicates the value is lower than lower limit of quantitation; n/d indicates the value is below assay detection limit.

[c] Data are presented as means ± standard errors from the mean. For germ-free controls, before gavage and colonized 1 week, n = 10 mice per group. For colonized 2 weeks and colonized 2 weeks + antibiotic water, n = 5 mice per group.

**Supplementary Table 3. Metabolite levels in *C. sporogenes* WT mono-colonized mouse urine samples**

| Metabolites (μM/mM Creatinine) | LLOQ[a] (μM) | Germ-free control[b,c] | Before gavage[b,c] | Colonized 1 week[b,c] | Colonized 2 weeks[b,c] | Colonized 2 weeks + antibiotics[b,c] |
|---|---|---|---|---|---|---|
| 5-Aminovalerate (5-AVA) | 9.28 | 3.51 ± 4.05 | 3.30 ± 2.09 | 350 ± 229 | 358 ± 115 | 3.69 ± 1.09 |
| Acetate | 594 | 529 ± 358 | 154 ± 102 | 175 ± 72.2 | 124 ± 60.1 | 62.1 ± 19.6 |
| Butyrate | 18.6 | <LLOQ | <LLOQ | <LLOQ | <LLOQ | <LLOQ |
| Hippurate | 18.6 | 22.7 ± 9.86 | 44.4 ± 9.46 | 1753 ± 465 | 1445 ± 198 | 42.2 ± 10.3 |
| 4-Hydroxyhippurate | 9.28 | 24.1 ± 6.08 | 28.0 ± 3.19 | 114 ± 27.5 | 88.5 ± 24.0 | 31.5 ± 5.35 |
| 4-Hydroxyphenylacetate (4-OH-PAA) | 9.28 | 18.3 ± 5.21 | 15.7 ± 4.53 | 22.2 ± 3.52 | 21.3 ± 2.50 | 3.55 ± 1.38 |
| 4-Hydroxyphenylacetylglycine (4-OH-PAG) | 2.32 | <LLOQ | 0.82 ± 0.19 | 0.75 ± 0.31 | 0.71 ± 0.13 | 0.58 ± 0.03 |
| 4-Hydroxyphenylpropionate (4-OH-PPA) | 4.64 | <LLOQ | 1.40 ± 0.60 | 187 ± 75.6 | 129 ± 17.4 | <LLOQ |
| 4-Hydroxyphenylpropionylglycine (4-OH-PPG) | 4.64 | <LLOQ | 1.18 ± 0.25 | 96.4 ± 34.6 | 89.0 ± 2.50 | 1.26 ± 0.15 |
| Indoleacetate (IAA) | 4.64 | <LLOQ | 1.50 ± 0.48 | 1.24 ± 0.37 | 1.39 ± 0.41 | 1.24 ± 0.30 |
| Indolepropionate (IPA) | 4.64 | n/d | n/d | n/d | n/d | n/d |
| Indolepropionylglycine (IPGly) | 4.64 | n/d | n/d | 73.2 ± 21.8 | 66.4 ± 21.9 | n/d |
| Isobutyrate | 9.28 | <LLOQ | 0.74 ± 0.33 | <LLOQ | <LLOQ | <LLOQ |
| Isocaproate | 9.28 | n/d | n/d | n/d | n/d | n/d |
| Isovalerate | 9.28 | <LLOQ | 1.32 ± 0.73 | <LLOQ | <LLOQ | 1.48 ± 0.71 |
| 2-Methylbutyrate | 18.6 | n/d | <LLOQ | <LLOQ | <LLOQ | <LLOQ |
| Phenylacetate (PAA) | 4.64 | 2.75 ± 1.01 | 2.96 ± 1.40 | 2.16 ± 0.95 | 3.20 ± 0.62 | 2.52 ± 1.14 |
| Phenylacetylglycine (PAG) | 4.64 | 63.7 ± 6.64 | 68.5 ± 8.54 | 84.1 ± 7.35 | 65.1 ± 6.30 | 69.4 ± 3.34 |
| Phenylpropionate (PPA) | 4.64 | n/d | n/d | n/d | n/d | n/d |
| Phenylpropionylglycine (PPG) | 4.64 | n/d | <LLOQ | 11.1 ± 3.22 | 8.46 ± 2.18 | <LLOQ |
| Propionate | 37.1 | <LLOQ | 3.04 ± 0.62 | 4.30 ± 1.54 | 2.91 ± 1.06 | 2.08 ± 0.86 |
| Tryptamine | 2.32 | <LLOQ | 0.22 ± 0.09 | <LLOQ | <LLOQ | <LLOQ |

[a] LLOQ is the lower limit of quantification for analytes prior to being divided by creatinine which was done to normalize values. The range of creatinine were 0.91 to 2.41 mM among these samples, and the LLOQ of creatinine was 0.018 mM.

[b] <LLOQ indicates the value is lower than lower limit of quantitation; n/d indicates the value is below assay detection limit.

[c] Data are presented as means ± standard errors from the mean. For germ-free controls, before gavage and colonized 1 week, n = 10 mice per group. For colonized 2 weeks and colonized 2 weeks + antibiotic water, n = 5 mice per group.

**Supplementary Table 4. Metabolite levels in *C. sporogenes* WT mono-colonized mouse cecal samples**

| Metabolites (nmol/mg)[a] | LLOQ (nmol/mg)[b] | Germ-free control[c,d] | Colonized 2 weeks[c,d] | Colonized 2 weeks + antibiotic water[c,d] |
|---|---|---|---|---|
| 5-Aminovalerate (5-AVA) | 0.019 | n/d | 9.02 ± 1.02 | n/d |
| Acetate | 0.30 | <LLOQ | 2.40 ± 0.28 | <LLOQ |
| Butyrate | 0.019 | <LLOQ | <LLOQ | <LLOQ |
| Hippurate | 0.005 | n/d | <LLOQ | n/d |
| 4-Hydroxyhippurate | 0.009 | n/d | n/d | n/d |
| 4-Hydroxyphenylacetate (4-OH-PAA) | 0.005 | <LLOQ | <LLOQ | <LLOQ |
| 4-Hydroxyphenylacetylglycine (4-OH-PAG) | 0.005 | n/d | n/d | n/d |
| 4-Hydroxyphenylpropionate (4-OH-PPA) | 0.009 | n/d | 0.72 ± 0.09 | n/d |
| 4-Hydroxyphenylpropionylglycine (4-OH-PPG) | 0.005 | n/d | n/d | n/d |
| Indoleacetate (IAA) | 0.005 | n/d | n/d | n/d |
| Indolepropionate (IPA) | 0.009 | n/d | 0.06 ± 0.01 | n/d |
| Isobutyrate | 0.009 | <LLOQ | 0.42 ± 0.07 | <LLOQ |
| Isocaproate | 0.009 | n/d | 0.02 ± 0.005 | n/d |
| Isovalerate | 0.019 | <LLOQ | 0.42 ± 0.07 | <LLOQ |
| 2-Methylbutyrate | 0.019 | n/d | 0.42 ± 0.06 | n/d |
| Phenylacetate (PAA) | 0.005 | <LLOQ | <LLOQ | <LLOQ |
| Phenylacetylglycine (PAG) | 0.005 | n/d | n/d | n/d |
| Phenylpropionate (PPA) | 0.005 | n/d | 0.21 ± 0.04 | n/d |
| Phenylpropionylglycine (PPG) | 0.002 | n/d | n/d | n/d |
| Propionate | 0.048 | <LLOQ | 0.22 ± 0.04 | <LLOQ |
| Tryptamine | 0.002 | n/d | <LLOQ | n/d |

[a] Data reported in the table were normalized by wet weight of feces (40 ± 4 mg).

[b] LLOQ is the lower limit of quantitation for the assay.

[c] <LLOQ indicates the value is lower than lower limit of quantitation; n/d indicates the value is below assay detection limit.

[d] Data are presented as means ± standard errors from the mean. For germ-free controls, n = 10 mice per group. For colonized 2 weeks and colonized 2 weeks + antibiotic water, n = 5 mice per group.

**Supplementary Table 5. Metabolite levels in human plasma samples**

| Blood Donor ID | 5-aminovalerate[a] (μM) (LLOQ 0.20 μM) | Phenylpropionate[a] (μM) (LLOQ 0.098 μM) | 3-(4-hydroxyphenyl)propionate[a] (μM) (LLOQ 0.098 μM) | Indolepropionate[a] (μM) (LLOQ 0.20 μM) |
|---|---|---|---|---|
| H-01 | 0.39 | 0.12 | n/d | 0.59 |
| H-02 | 0.74 | 0.27 | n/d | 0.79 |
| H-03 | 0.31 | 0.06 | n/d | 0.57 |
| H-04 | 0.15 | 0.06 | n/d | 0.23 |
| H-05 | 0.23 | 0.10 | 0.04 | 0.56 |
| H-06 | 0.50 | 0.06 | 0.03 | 0.40 |
| H-07 | 0.17 | 0.19 | n/d | 1.03 |
| H-08 | n/d | 0.08 | n/d | 0.25 |
| H-09 | 0.46 | 0.29 | n/d | 0.51 |
| H-10 | 0.50 | 0.08 | n/d | 0.28 |
| H-11 | 0.50 | 0.05 | 0.03 | 1.37 |
| H-12 | 0.21 | 0.17 | n/d | 1.40 |
| H-13 | 0.23 | 0.06 | 0.05 | 0.50 |
| H-14 | 0.37 | 0.32 | 0.04 | 1.50 |
| H-15 | 0.39 | 0.05 | 0.03 | 0.60 |
| H-16 | 0.17 | 0.07 | 0.03 | 0.75 |
| H-17 | 0.39 | 0.52 | n/d | 1.19 |
| H-18 | 0.55 | 0.36 | n/d | 1.09 |
| H-19 | 0.25 | 0.22 | n/d | 0.68 |
| H-20 | 0.14 | 0.20 | 0.02 | 1.55 |
| H-21 | 0.14 | 0.06 | 0.04 | 5.09 |
| H-22 | 0.81 | 0.23 | 0.03 | 0.67 |
| H-23 | 0.15 | 1.63 | 0.07 | 4.86 |
| H-24 | n/d | 0.12 | n/d | 1.26 |
| H-25 | 0.12 | 0.63 | n/d | 0.86 |
| H-26 | 0.33 | 0.70 | n/d | 2.11 |
| H-27 | 0.14 | 0.09 | 0.03 | 1.15 |
| H-28 | 0.14 | 0.35 | n/d | 1.78 |
| H-29 | 0.27 | 0.54 | n/d | 0.73 |
| H-30 | 0.11 | 0.44 | n/d | 0.89 |
| H-31 | 0.30 | 0.49 | n/d | 0.78 |
| H-32 | 0.59 | 1.07 | n/d | 3.85 |
| H-33 | 0.74 | 0.22 | n/d | 0.35 |
| H-34 | n/d | 0.26 | n/d | 1.53 |
| H-35 | 0.13 | 0.27 | 0.03 | 0.49 |
| H-36 | n/d | n/d | 0.07 | 0.43 |
| H-37 | 0.10 | 0.28 | n/d | 1.22 |
| H-38 | 0.14 | n/d | 0.03 | n/d |
| H-39 | 0.51 | 2.57 | 0.07 | 112.60 |
| H-40 | n/d | 0.16 | 0.19 | 3.00 |

| | | | | |
|---|---|---|---|---|
| H-41 | 0.11 | 0.16 | n/d | 0.51 |
| H-42 | 0.15 | n/d | 0.11 | 1.17 |
| H-43 | 0.21 | 0.45 | n/d | 1.09 |
| H-44 | n/d | 0.34 | n/d | 0.47 |
| H-45 | n/d | n/d | 0.08 | 1.47 |
| H-46 | 0.21 | 0.04 | 0.03 | 0.52 |
| H-47 | 0.29 | 0.43 | n/d | 2.85 |
| H-48 | n/d | 0.58 | n/d | 3.45 |
| H-49 | 0.33 | 0.28 | n/d | 1.28 |
| H-51 | n/d | 6.32 | 0.05 | 5.66 |
| H-52 | 0.10 | 0.05 | n/d | 0.37 |
| H-53 | 0.19 | n/d | 0.04 | 0.16 |
| H-54 | 0.17 | 0.70 | n/d | 7.34 |
| H-55 | 0.17 | n/d | 0.04 | 0.77 |
| H-56 | n/d | 1.58 | n/d | 2.63 |
| H-57 | 0.19 | 0.27 | n/d | 1.75 |
| H-58 | 0.17 | 0.90 | n/d | 1.02 |
| H-59 | 0.10 | 0.42 | n/d | 1.69 |
| H-60 | 0.78 | 0.24 | n/d | 1.53 |
| H-61 | 0.31 | 0.22 | 0.03 | 0.63 |
| H-62 | 0.17 | 0.08 | 0.11 | 2.72 |
| H-63 | 0.24 | 0.61 | 0.04 | 1.61 |
| H-64 | n/d | 0.08 | n/d | 0.95 |
| H-65 | 0.16 | 0.45 | n/d | 9.27 |
| H-66 | n/d | 1.03 | n/d | 1.65 |
| H-67 | 0.22 | 0.40 | n/d | 1.65 |
| H-68 | 0.15 | 0.15 | n/d | 0.46 |
| H-69 | 0.35 | 1.06 | n/d | 1.19 |
| H-70 | n/d | 0.08 | n/d | 0.55 |
| H-71 | 0.22 | 0.91 | 0.03 | 0.47 |
| H-72 | 0.13 | 0.06 | 0.08 | 2.23 |
| H-73 | 0.15 | 0.85 | n/d | 0.71 |
| H-74 | 0.52 | 0.33 | 0.04 | 2.09 |
| H-75 | 0.59 | 0.60 | n/d | 1.33 |
| H-76 | 0.21 | 0.08 | n/d | 1.08 |
| H-77 | 0.13 | 0.43 | 0.03 | 1.85 |
| H-78 | 0.23 | 0.27 | 0.04 | 0.94 |
| H-79 | 0.24 | 1.25 | 0.03 | 0.90 |
| H-80 | 0.42 | 0.28 | n/d | 0.99 |
| H-81 | 0.19 | 0.58 | 0.03 | 0.51 |
| H-82 | 0.15 | 0.93 | 0.05 | 1.02 |
| H-83 | 0.10 | 0.60 | n/d | 1.13 |
| H-84 | 0.11 | 0.17 | n/d | 2.47 |
| H-85 | n/d | 0.87 | n/d | 3.63 |
| H-86 | 0.23 | 0.27 | n/d | 1.31 |

| | | | | |
|---|---|---|---|---|
| H-87 | 0.17 | 0.15 | 0.05 | 0.80 |
| H-88 | 0.13 | 0.28 | n/d | 2.04 |
| H-89 | 0.09 | 0.63 | 0.03 | 1.74 |
| H-90 | 0.21 | 0.09 | 0.06 | 0.62 |
| H-91 | 0.12 | 0.24 | n/d | 1.00 |
| H-92 | n/d | 0.23 | n/d | 0.50 |
| H-93 | n/d | 0.20 | n/d | 0.46 |
| H-94 | 0.10 | 0.90 | n/d | 5.25 |
| H-95 | 0.14 | 0.26 | n/d | 0.69 |
| H-96 | 0.22 | n/d | 0.03 | 3.21 |
| H-97 | n/d | 0.41 | n/d | 3.17 |
| H-98 | 0.24 | 0.24 | n/d | 1.44 |
| H-99 | n/d | 0.19 | n/d | 2.00 |
| H-100 | 0.17 | 0.30 | 0.12 | 4.92 |

[a] n/d indicates the value is below assay detection limit.

**Supplementary Table 7. Estimates of ATP levels during Stickland metabolism of Val/Phe.**

| Considerations | Oxidative Pathway (Val) | Reductive Pathway (Phe) |
|---|---|---|
| Stoichiometry for redox balance | 1 mole Val | 2 moles Phe |
| NADH reducing equivalents (total) | 2 NADH produced | 2 NADH produced<br>4 NADH consumed<br>Net: 2 NADH consumed |
| ATP from substrate level phosphorylation | 1 ATP | |
| Ferredoxin reduced (2 e- reduction) | 1 mole $Fd^{red}$ | 2 moles $Fd^{red}$ |
| Protons translocated through Rnf complex (assuming 2 protons per mole $Fd^{red}$)[a] | 2 protons | 4 protons |
| ATP from ETP (assuming 4 protons per mole ATP)[a] | 0.5 ATP | 1 ATP |
| Total ATP | 1.5 ATP | 1 ATP |
| Percentage of total ATP | 60% | 40% |

[a] Estimates are from Buckel and Thauer[17].

**Supplementary Table 8. Primers used in this study.**

Sequencing primers for ClosTron mutants

| Gene (Locus ID) | Targeted Region[a] | Sequencing Primers (5′→3′)[b,c] |
|---|---|---|
| *rnfB* (CLOSPO_00568) | 422s | rnfBF: TTTACCAGGAGCTAACTGTG |
| | | rnfBR: TGGCAAATATCTTTAACAGC |
| *rnfE* (CLOSPO_00570) | 61s | rnfEF: TTTAGAAGTTGTTAAGACAGC |
| | | rnfBR: TATTGAAGATACTGGACCAT |

Q-PCR primers

| Primer | Orientation | Sequence (5′→3′)[c] |
|---|---|---|
| *Clostridium sporogenes* total | Forward | TTGGCTCTGCACCGGGAATC |
| | Reverse | CTGCAAACGCCGTCCCTCTT |
| *C. sporogenes rnfB* mutant | Forward | AGAGCAACCCTAGTGTTCGGTGA |
| | Reverse | TTGAAAGCCATGCGTCTGACATCT |
| *Bifidobacterium breve UCC2003* | Forward | ACAGTCTTGCGCGTAAAGCT |
| | Reverse | GCACGGGCCTCAAACGTATA |
| *Bacteroides thetaiotaomicron VPI-5482* | Forward | AGCCGAGAACTCCATGTCTTT |
| | Reverse | TGCTACCAGCGATTGCATAGA |
| *Bacteroides vulgatus ATCC 8482* | Forward | CGTACGAGGATGGACGCATT |
| | Reverse | AGTGCACCCGAACCCAATAC |
| *Clostridium scindens ATCC 35704* | Forward | TTCAGGAGAAATGCCAGCCAGT |
| | Reverse | GTTCTGCCACGGTCGTTAGC |
| *Eubacterium rectale ATCC 33656* | Forward | ACGCCCACTCACGGATCTGA |
| | Reverse | GCCAAACAGCCCCGACATCA |
| *Edwardsiella tarda ATCC 23685* | Forward | ATGTGCCTGTTGATTTGCCC |
| | Reverse | CGCAGGTGGGTATGGTGATT |

[a] Targeting regions were designed using the Intron design tool on the ClosTron website (http://www.clostron.com/clostron2.php). The regions listed indicate the nucleotide position within the gene where the intron is inserted and the a/s designation indicates whether the intron integrates in the antisense or sense strand.

[b] Primers were designed to amplify the region containing the expected intron after integration.

[c] Primers were synthesized by Integrated DNA Technologies (Coralville, IA).

### Supplementary Figures

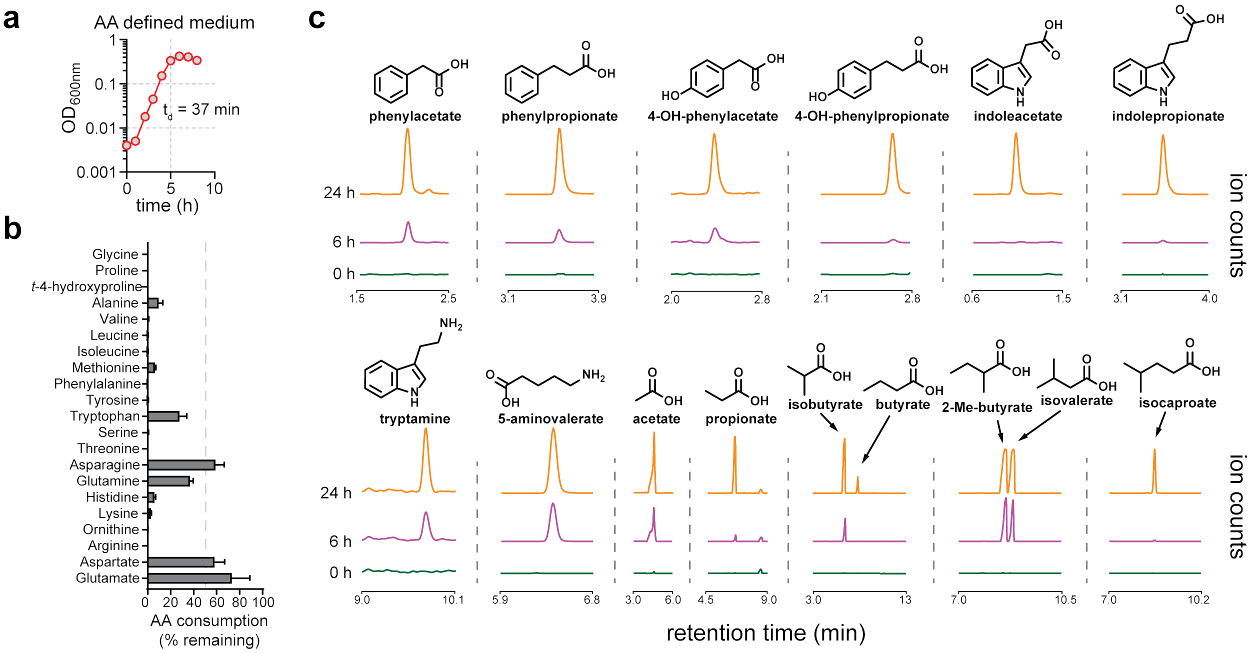


**Extended data Fig. 1. *Clostridium sporogenes* produces fifteen small molecules during amino acid metabolism.** *C. sporogenes* A) grows rapidly in amino acid defined, B) depletes amino acids during growth, and C) produces fifteen metabolites. All experiments were repeated independently 3 times. For B, amino acid levels are plotted as means +/- standard deviations from the mean. For A and C, representative data are shown.

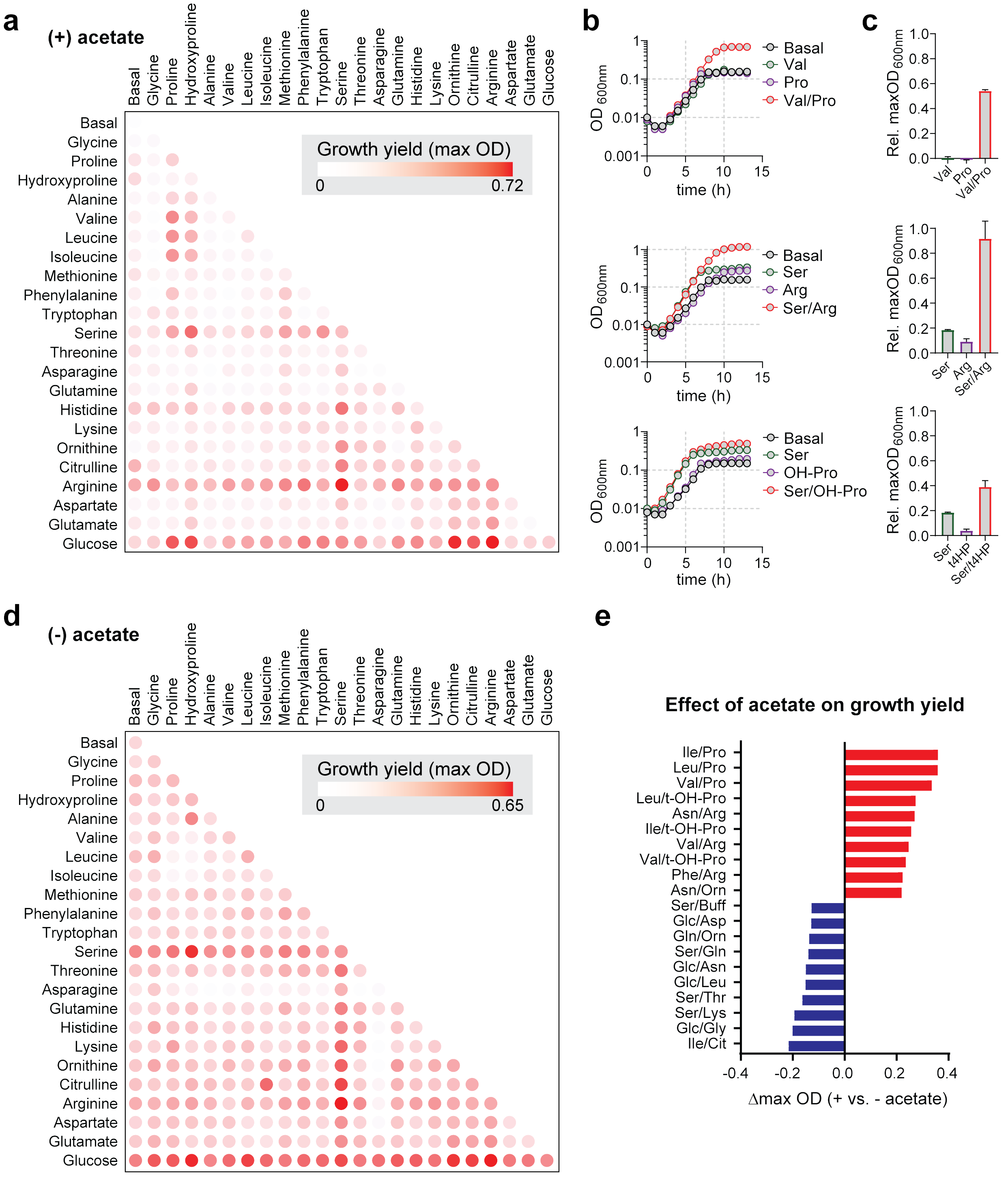


**Extended data Fig. 2. Growth-based assay reveals coupled amino acid metabolism in *C. sporogenes*.** A) *C. sporogenes* was grown in basal medium containing 10 essential amino acids (Gly, Val, Leu, Ile, His, Met, Arg, Phe, Tyr, Trp; 1 mM each) supplemented with acetate (40 mM) and indicated substrates (25 mM) for 48 h and the maximum optical density at 600nm was recorded using a microplate spectrophotometer. B-C) Hit validation in batch culture reveals synergism among specific amino acid pairs. D) Experiments were performed as in (A), except that no supplemental acetate was provided. E) Effect of acetate supplementation on maximum growth yields from substrate combinations. The change in maximum optical density represents the maximum OD during growth without acetate subtracted from maximum OD during growth with acetate. The top ten pairs of substrates where acetate promoted growth are shown in red. The top ten pairs of substrates where acetate diminished growth are shown in blue. In A, the growth screen was performed two times independently with technical duplicates. In D, the growth screen was performed once with technical duplicates. For B-C, experiments were repeated independently three times. For A-D, representative data are shown.

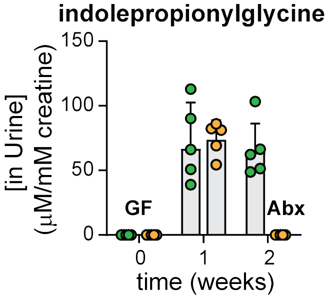


**Extended data Fig. 3. Indolepropionylglycine (IPGly) is the excreted form of indolepropionate in urine of mice mono-colonized with *C. sporogenes*.** Indolepropionylglycine was quantified in urine of mice described in **Fig. 1b** using LC-MS and normalized to urine creatinine levels. Means and standard deviations for urine IPGly are reported in **Supplementary Table 3**.

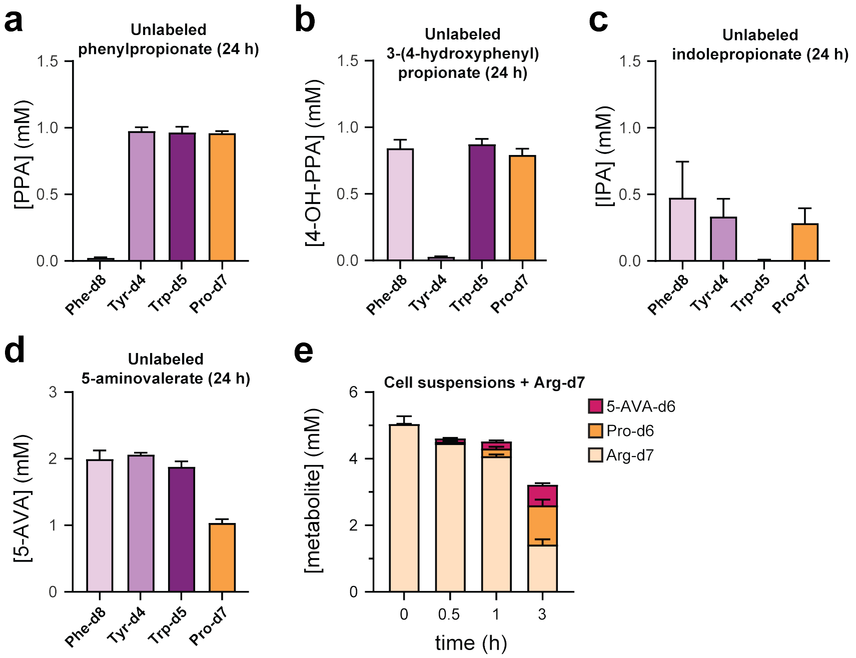


**Extended data Fig. 4. Stickland metabolites do not arise from *de novo* biosynthetic pathways.** A-D) Stable isotope tracing. *C. sporogenes* was cultured in a synthetic medium containing 20 amino acids where Phe, Tyr, Trp, and Pro were individually substituted by their deuterium isotopologues. Cell-free supernatants (n = 3 per group) were collected at t = 0, 6, 24 h and metabolites were detected by LC-MS. A-C) For phenylpropionate, 3-(4-hydroxyphenyl)propionate, and indolepropionate, no unlabeled products were detected when isotopically labeled amino acids were provided. D) Unlabeled 5-aminovalerate levels were reduced when isotopically labeled Pro was supplied, but ~1 mM 5-aminovalerate was still detected suggesting another source exists for its production. E) Stable isotope tracing of *C. sporogenes* cell suspensions incubated with stable isotopically labeled arginine (Arg-d7). When cells were incubated with Arg-d7, labeled proline and 5-aminovalerate were detected, suggesting that arginine is a substrate for 5-aminovalerate production.

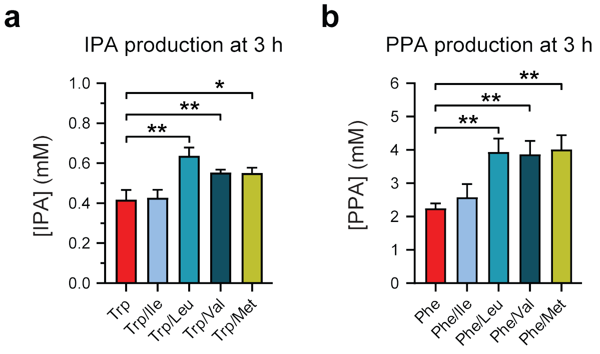


**Extended data Fig. 5. Indolepropionate (IPA) production is stimulated by incubation with oxidative Stickland amino acids.** A) *C. sporogenes* cell suspensions were incubated with stable isotopically labeled tryptophan (Trp-d5) alone or in combination with oxidative Stickland amino acids (Ile, Leu, Val, or Met), then labeled IPA-d5 was measured by LC-MS. IPA-d5 levels were stimulated by addition of Leu, Val, or Met. B) *C. sporogenes* cell suspensions were incubated with stable isotopically labeled phenylalanine (Phe-d8) alone or in combination with oxidative Stickland amino acids (Ile, Leu, Val, or Met), then labeled PPA-d6 was measured by LC-MS. PPA-d6 levels were stimulated by addition of Leu, Val, or Met. Experiments were performed in triplicate and data are reported as means ± standard deviations. *, $p<0.05$; **, $p<0.01$

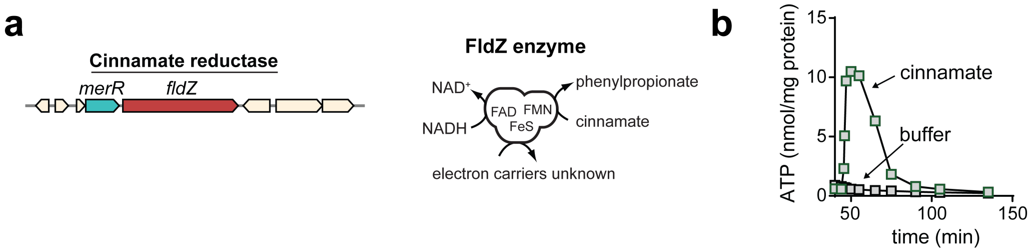


**Extended data Fig. 6. Cinnamate reduction is coupled to ATP formation in *C. sporogenes*.** A) The cinnamate reductase gene encodes a unique FAD/[FeS]/FMN containing enzyme with separate binding sites for NADH, cinnamate, and the artificial electron carrier methylviologen, however the natural electron carriers remain unknown. Cinnamate can be reduced either by NADH or methylviologen. B) Resting cell suspensions of *C. sporogenes* accumulate ATP after being incubated with cinnamate. Buffer control is shown. For B, experiments were repeated independently three times and representative data are shown.

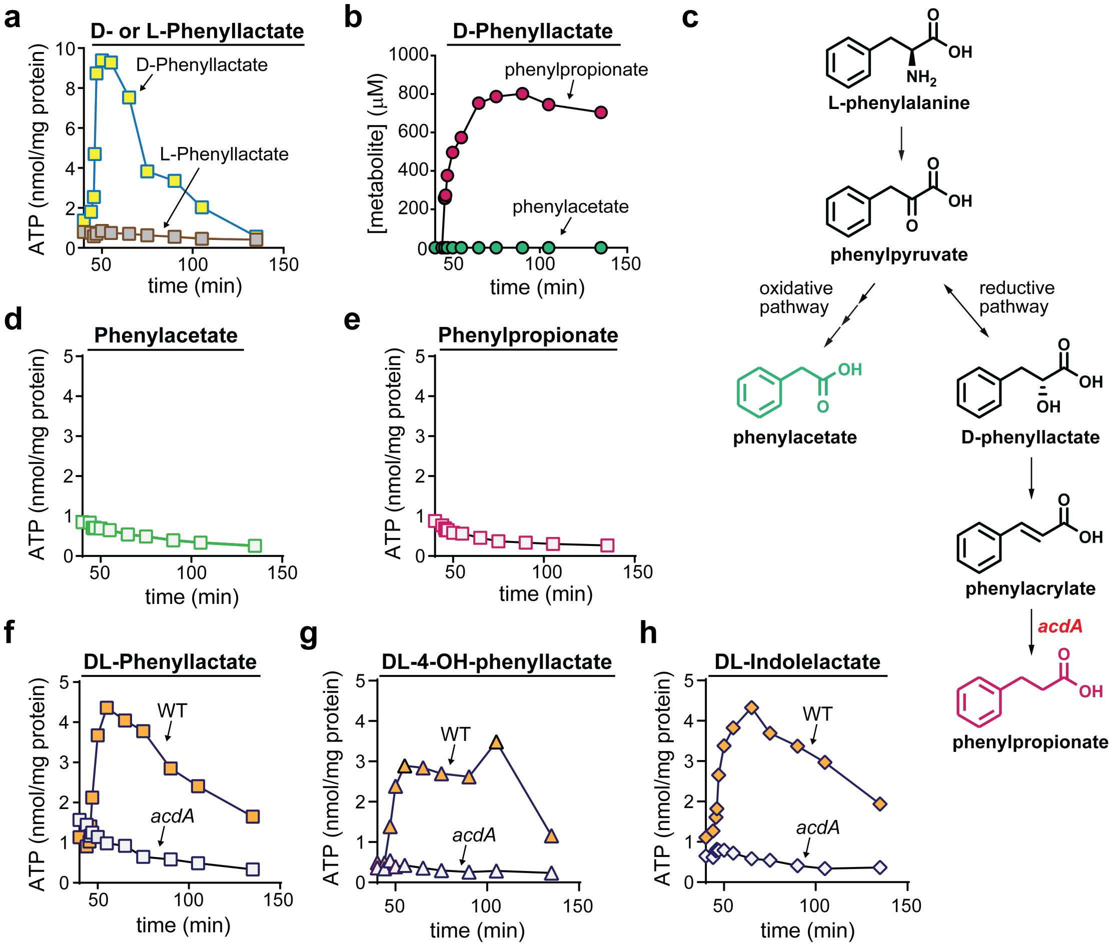


**Extended data Fig. 7. Reductive metabolism of D-phenyllactate is coupled to ATP formation involving *acdA*.** A) D- or L-phenyllactate was added (1 mM) to resting cell suspensions of *C. sporogenes*, and ATP levels were measured at indicated time points using a luciferase-based assay. B) Reductive (PPA) and oxidative (PAA) pathway end products were measured by tandem mass spectrometry during D-phenyllactate metabolism. PPA, phenylpropionate; PAA, phenylacetate. C) Pathways for oxidative and reductive metabolism of phenylalanine showing the position of AcdA in the pathway. D-E) Phenylacetate (D) or phenylpropionate (E) was added (1 mM) to resting cell suspensions of *C. sporogenes*, and ATP levels were measured at indicated time points using a luciferase-based assay. F-H) DL-phenyllactate (F), DL-3-(4-hydroxyphenyl)lactate (G), or DL-indolelactate (H) was added (1 mM) to resting cell suspensions of WT or *acdA* mutant *C. sporogenes*, and ATP levels were measured at indicated time points using a luciferase-based assay. For A-B, D-H, experiments were repeated independently three times and representative data are shown.

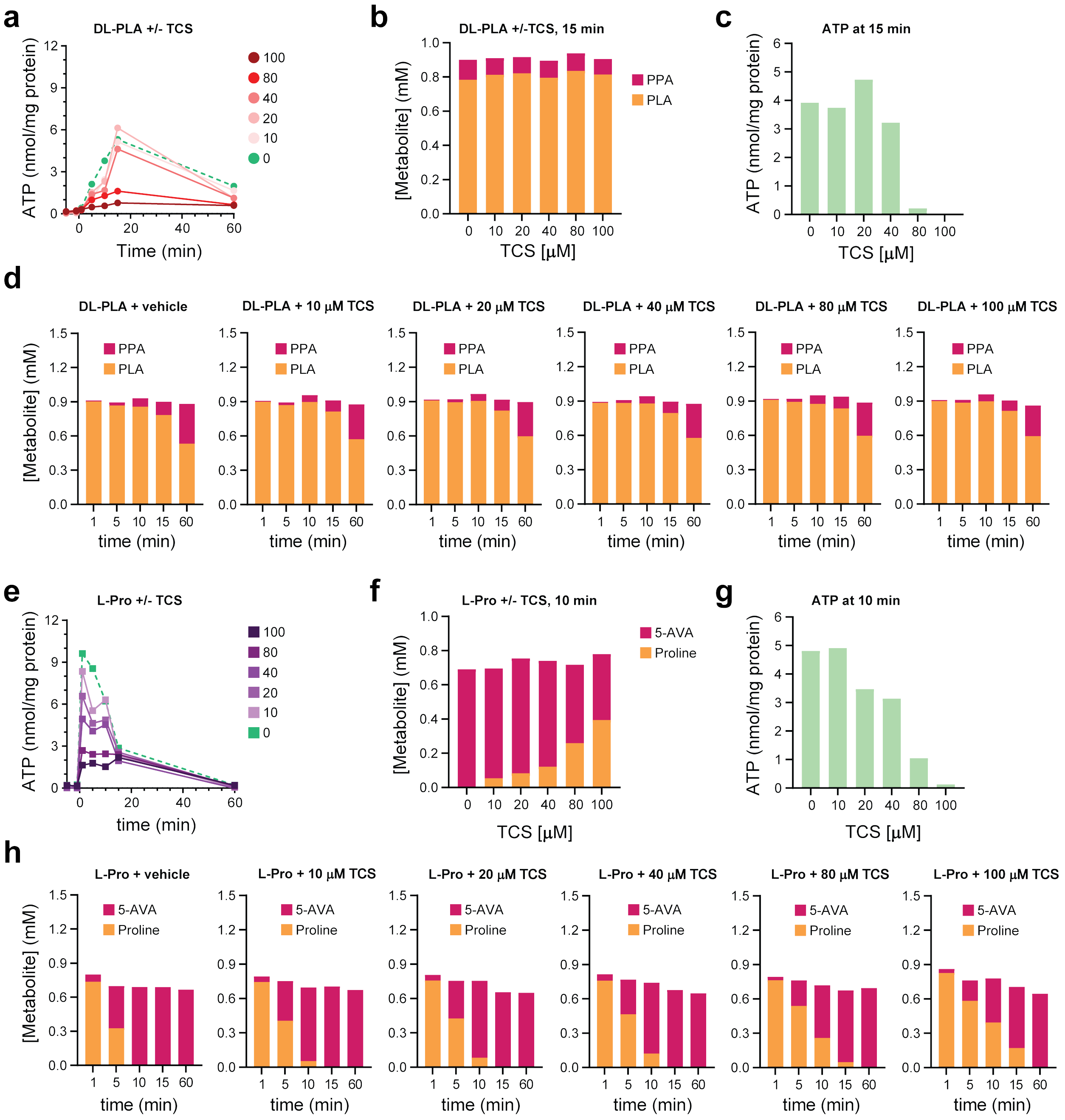


**Extended data Fig. 8. The protonophore, 3,3′,4′,5-tetrachlorosalicylanilide (TCS), uncouples reductive Stickland metabolism from ATP formation.** Resting cell suspensions of *C. sporogenes* were preincubated with ethanol (vehicle) or varying concentrations of TCS for 30 min. Then substrates (DL-phenyllactate (DL-PLA) or proline (Pro)) were added, and aliquots were taken at different time-points and quenched in DMSO. Total cellular ATP was quantified using a luciferase-based assay and normalized to total cellular protein and substrates/metabolites were quantified using LC-MS as described in the methods. A) Dose dependent decrease in ATP formation from DL-phenyllactate with increasing concentrations of TCS. B) Conversion of DL-phenyllactate to phenylpropionate (PPA) at the 15 min timepoint remains constant with increasing TCS while (C) ATP production diminishes. D) Conversion of DL-phenyllactate to phenylpropionate (PPA) increases over time irrespective of TCS levels. E) Dose dependent decrease in ATP formation from Pro with increasing concentrations of TCS. F) Conversion of Pro to 5-aminovalerate (5-AVA) at the 15 min timepoint decreases but is not completely blocked with increasing TCS while (G) ATP production diminishes becoming negligible at 100 μM TCS. H) Conversion of Pro to 5-aminovalerage (5-AVA) increases over time, and while the rate of conversion decreases with increasing TCS levels, proline conversion to 5-aminovalerate is complete by the 60 min timepoint.

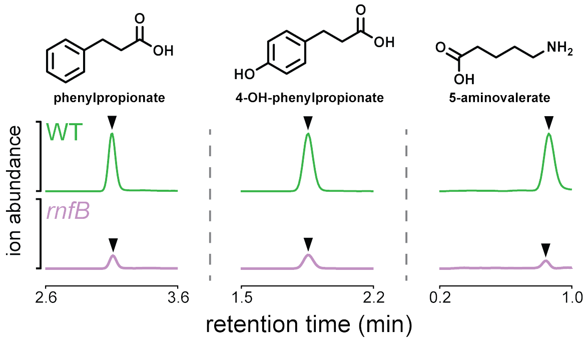


**Extended data Fig. 9. The *rnfB* mutant produces less phenylpropionate, 3-(4-hydroxyphenyl)propionate, and 5-aminovalerate during growth in defined medium.** Wild-type and *rnfB* mutant *C. sporogenes* were cultured in SACC medium with 10 amino acids and no glucose for 15 h, then metabolites were detected in culture supernatants by LC-MS.

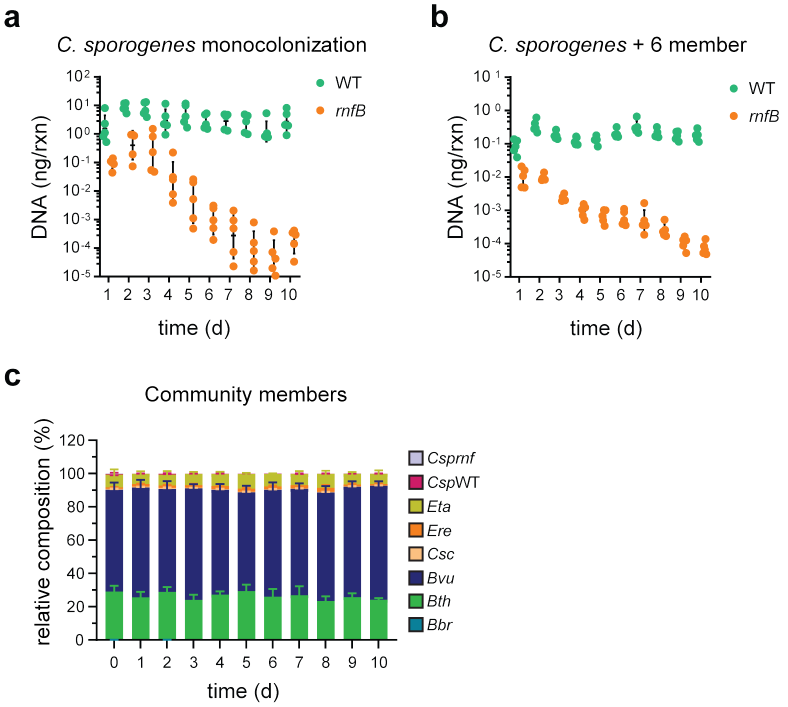


**Extended data Fig. 10. Levels of wild-type and *rnfB* mutant *C. sporogenes* in gnotobiotic competition experiments and stability of defined community members over time.** A) Germ-free mice were colonized by oral gavage of a 1:1 mixture of wild-type (WT) or *rnfB* mutant *C. sporogenes* and feces were collected daily for 10 days and genomic DNA was purified. Wild-type and *rnfB* mutant *C. sporogenes* were quantified by Q-PCR and plots show the amount of wild-type or *rnfB* mutant DNA per PCR reaction. B-C) Germ-free mice were colonized by a defined microbial consortium consisting of 6 bacteria (*Edwardsiella tarda* ATCC 23685 (Eta), *Eubacterium rectale* ATCC 33656 (Ere), *Clostridium scindens* ATCC 35704 (Csc), *Bacteroides vulgatus* ATCC 8482 (Bvu), *Bacteroides thetaiotaomicron* VPI-5482 (Bth), and *Bifidobacterium breve* UCC2003 (Bbr)). One week later, mice were administered by oral gavage a 1:1 mixture of wild-type or *rnfB* mutant *C. sporogenes* and feces were collected daily for 10 days and genomic DNA was purified. In B) wild-type and *rnfB* mutant *C. sporogenes* were quantified by Q-PCR and plots show the amount of wild-type or *rnfB* mutant DNA per PCR reaction. For C), taxa plots were generated using Q-PCR with organism specific primers for all community members over time.

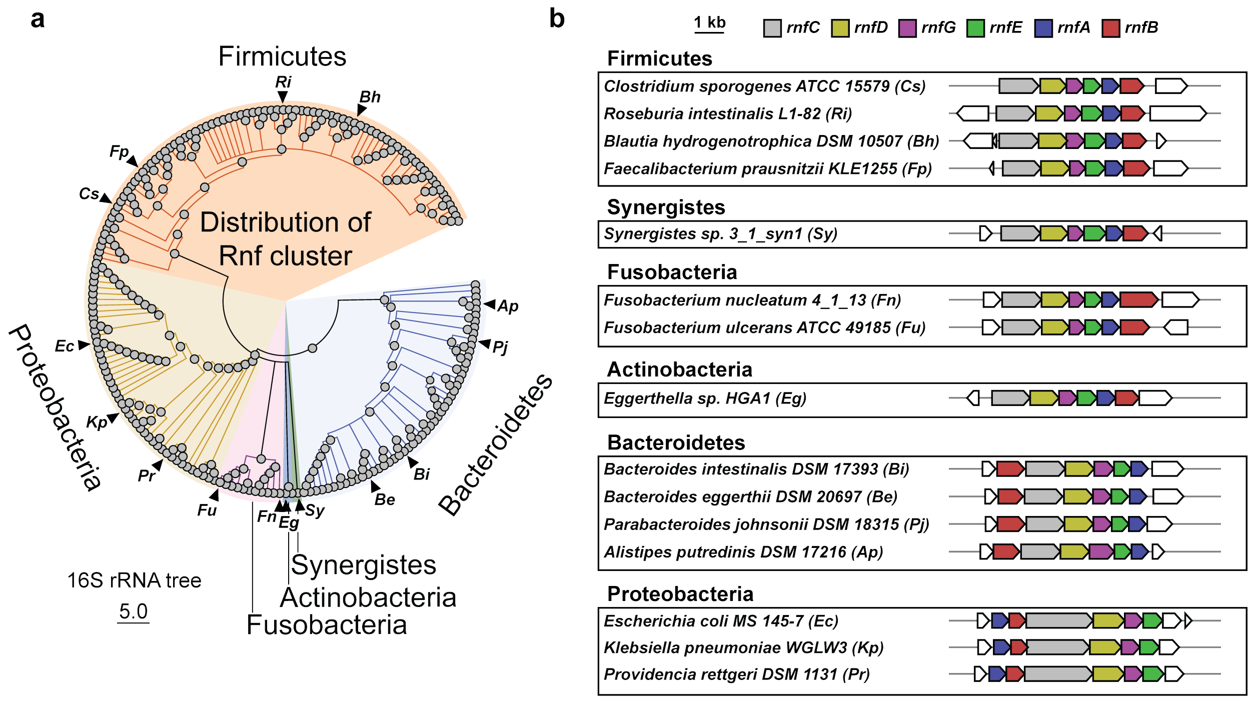


**Extended data Fig. 11. The Rnf gene cluster is widely distributed among reference genomes from the Human Microbiome Project.** A) 16S rRNA tree of organisms from the human microbiome reference genome collection determined to have Rnf gene clusters as determined by BLASTp and manual inspection of gene neighborhoods using MultiGeneBlast. B) Rnf gene clusters for select organisms (corresponding to triangles in panel A), grouped by phyla.

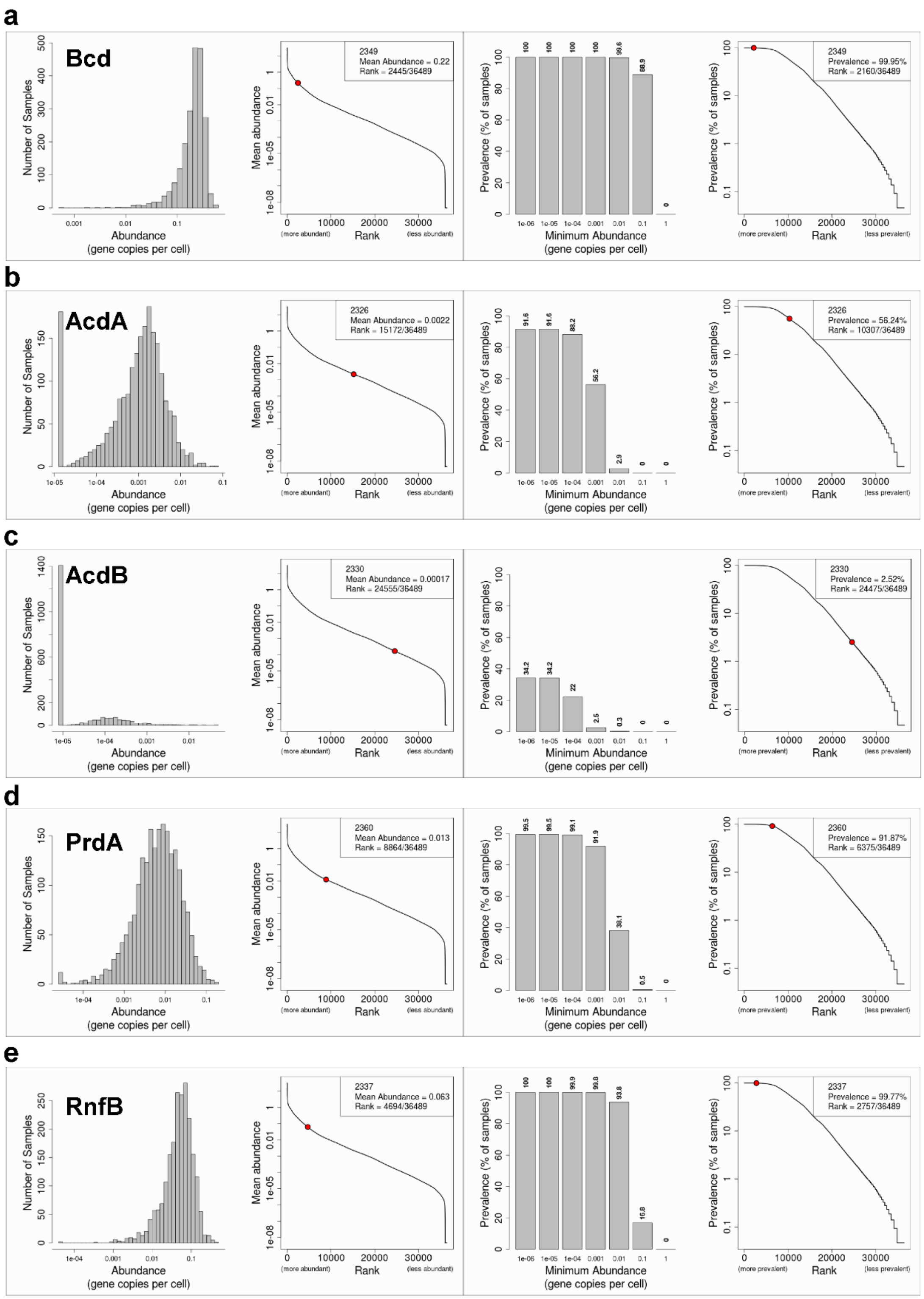


**Extended data Fig. 12. Abundance and prevalence of electron transfer proteins within human fecal metagenomic datasets.** *C. kluyveri* Bcd (EDK32509.1), *C. sporogenes* AcdA (EDU39257.1), AcdB (EDU36591.1), PrdA (EDU36353.1), or RnfB (EDU37753.1) were used as a query to search ~2,000 human microbiome datasets using the metaquery tool (http://metaquery.docpollard.org/). On the left, plots show the relative abundance of the query genes and on the right, plots show the prevalence of the query genes.

### Supplementary References


1 Wildenauer, F. X. & Winter, J. Fermentation of isoleucine and arginine by pure and syntrophic cultures of *Clostridium sporogenes*. *FEMS Microbiol Lett* **38**, 373-379 (1986).

2 Lovitt, R. W., Morris, J. G. & Kell, D. B. The growth and nutrition of *Clostridium sporogenes* NCIB 8053 in defined media. *J Appl Bacteriol* **62**, 71-80 (1987).

3 Lovitt, R. W., Kell, D. B. & Morris, J. G. The physiology of *Clostridium sporogenes* NCIB 8053 growing in defined media. *J Appl Bacteriol* **62**, 81-92 (1987).

4 Bouillaut, L., Self, W. T. & Sonenshein, A. L. Proline-dependent regulation of *Clostridium difficile* Stickland metabolism. *J Bacteriol* **195**, 844-854, doi:10.1128/JB.01492-12 (2013).

5 Jackson, S., Calos, M., Myers, A. & Self, W. T. Analysis of proline reduction in the nosocomial pathogen *Clostridium difficile*. *J Bacteriol* **188**, 8487-8495, doi:10.1128/JB.01370-06 (2006).

6 Hofmeister, A. E., Grabowski, R., Linder, D. & Buckel, W. L-serine and L-threonine dehydratase from *Clostridium propionicum*. Two enzymes with different prosthetic groups. *Eur J Biochem* **215**, 341-349, doi:10.1111/j.1432-1033.1993.tb18040.x (1993).

7 Xu, X. L. & Grant, G. A. Identification and characterization of two new types of bacterial L-serine dehydratases and assessment of the function of the ACT domain. *Arch Biochem Biophys* **540**, 62-69, doi:10.1016/j.abb.2013.10.009 (2013).

8 Zhang, X., El-Hajj, Z. W. & Newman, E. Deficiency in L-serine deaminase interferes with one-carbon metabolism and cell wall synthesis in *Escherichia coli* K-12. *J Bacteriol* **192**, 5515-5525, doi:10.1128/JB.00748-10 (2010).

9 Leach, S., Harvey, P. & Wali, R. Changes with growth rate in the membrane lipid composition of and amino acid utilization by continuous cultures of *Campylobacter jejuni*. *J Appl Microbiol* **82**, 631-640, doi:10.1111/j.1365-2672.1997.tb02873.x (1997).

10 Velayudhan, J., Jones, M. A., Barrow, P. A. & Kelly, D. J. L-serine catabolism via an oxygen-labile L-serine dehydratase is essential for colonization of the avian gut by *Campylobacter jejuni*. *Infect Immun* **72**, 260-268, doi:10.1128/iai.72.1.260-268.2004 (2004).

11 Giesel, H. & Simon, H. On the occurrence of enoate reductase and 2-oxo-carboxylate reductase in clostridia and some observations on the amino acid fermentation by *Peptostreptococcus anaerobius*. *Arch Microbiol* **135**, 51-57 (1983).

12 Dickert, S., Pierik, A. J. & Buckel, W. Molecular characterization of phenyllactate dehydratase and its initiator from *Clostridium sporogenes*. *Mol Microbiol* **44**, 49-60, doi:10.1046/j.1365-2958.2002.02867.x (2002).

13 Mordaka, P. M., Hall, S. J., Minton, N. & Stephens, G. Recombinant expression and characterisation of the oxygen-sensitive 2-enoate reductase from *Clostridium sporogenes*. *Microbiology* **164**, 122-132, doi:10.1099/mic.0.000568 (2018).

14 Dodd, D. *et al.* A gut bacterial pathway metabolizes aromatic amino acids into nine circulating metabolites. *Nature* **551**, 648-652 (2017).
15 Bader, J. & Simon, H. ATP formation is coupled to the hydrogenation of 2-enoates in *Clostridium sporogenes*. *FEMS Microbiol Lett* **20**, 171-175 (1983).
16 Hess, V., Gonzalez, J. M., Parthasarathy, A., Buckel, W. & Muller, V. Caffeate respiration in the acetogenic bacterium *Acetobacterium woodii*: a coenzyme A loop saves energy for caffeate activation. *Appl Environ Microbiol* **79**, 1942-1947 (2013).
17 Buckel, W. & Thauer, R. K. Flavin-based electron bifurcation, ferredoxin, flavodoxin, and anaerobic respiration with protons (Ech) or NAD(+) (Rnf) as electron acceptors: A historical review. *Front Microbiol* **9**, 401 (2018).
